# A *de novo* CO_2_ Reductase Featuring a Cysteine-Ligated Cobalt Porphyrin Cofactor

**DOI:** 10.64898/2026.05.07.723500

**Authors:** Emily J. Radley, Alessia C. Andrews, Indrek Kalvet, Yunling Deng, Colin W. Levy, Mary Ortmayer, Derren J. Heyes, Clare F. Megarity, Reyes Núñez-Franco, Amy E. Hutton, Yi Lu, David Baker, Anthony P. Green

**Author notes:** These authors contributed equally (Emily J. Radley, Alessia C. Andrews).

## Abstract

Modern protein design methods based on deep learning allow generation of customized protein scaffolds with diverse geometries and functionalities. Here, we capitalize on these recent advances to develop hyper-thermostable *de novo* CO_2_ reductases featuring a cobalt porphyrin IX cofactor (CoPPIX). CoPPIX containing enzymes were assembled *in vivo* through media supplementation with cobalt salts and assessed for photocatalytic CO_2_ reductase activity. We identified two cysteine-ligated designs that exhibit high activity (>1000 turnovers at rates of up to 25 min^-1^) while suppressing competing hydrogen evolution pathways. A 2.1 Å crystal structure shows close agreement to the design model with the Co-Cys bond programmed as intended. This study showcases the power of computational protein design in developing artificial enzymes to activate challenging molecules such as CO_2_.

## Introduction

Rising CO_2_ levels in the atmosphere are a major driver of global warming. To facilitate the transition to net-zero, new catalytic technologies to transform CO_2_ into platform chemicals are needed. In this regard, a variety of homogeneous and heterogeneous catalysts have been developed for reducing CO_2_ to CO and other C1 products^1–10^. Although these systems can offer high catalytic rates, they commonly suffer from poor stability and/or compromised selectivity as they are unable to discriminate between competing CO_2_ and proton reduction pathways^11,12^. In contrast, enzymes have evolved in nature to transform CO_2_ with remarkable efficiency and selectivity^13–21^. Unfortunately, these natural systems are often only marginally stable and can be challenging to produce and characterize^22–24^. As a result, the structural features that underpin activation and transformation of CO_2_ in native enzymes are still not fully understood. One approach to overcome these limitations is to develop artificial metalloenzymes within small stable protein scaffolds that are amenable to engineering^25–27^. This approach can provide new insights into the factors governing selective CO_2_ conversion and lead to the development of biocatalysts with improved properties. As an example, we have recently developed artificial CO_2_ reductases by reconstituting myoglobin variants with a non-native cobalt protoporphyrin IX (CoPPIX) cofactor^28^. These artificial enzymes mediate photocatalytic conversion of CO_2_ to CO with enhanced activity and selectivity (CO formation vs H_2_ production) compared to small synthetic Co-porphyrin cofactors^27–31^.

In principle, these myoglobin-based systems could serve as starting templates for evolutionary optimization to deliver highly active, selective, and specialized CO_2_ reductases. However, establishing suitable high throughput directed evolution pipelines for this class of transformation presents considerable challenges linked to difficulties in performing high throughput quantitative analysis of gas phase products. In light of these constraints, we instead turned to modern protein design methods in an effort to generate artificial metalloenzymes with improved properties. Our labs have recently developed hyper-stable peroxidases and carbene transferases featuring a histidine-ligated Fe-porphyrin in a *de novo* α-helical solenoid repeat scaffold^32^. These studies provided insights into the factors leading to high-affinity porphyrin binding within *de novo* proteins. Armed with this knowledge, and with powerful deep learning-based methods now available for generating protein ligand complexes (e.g. RFdiffusion All-Atom^33^ and LigandMPNN^34^), we have the opportunity to create porphyrin-binding proteins with diverse active site geometries, cofactor coordination environments, and topologies that extend far beyond those found in nature. We set out to use these methods to generate a family of structurally diverse CoPPIX binding proteins and investigate their CO_2_ reductase activities.

## Results and Discussion

### *In vivo* assembly of a *de novo* CO_2_ reductase

As a starting template for engineering *de novo* CO_2_ reductases we selected dnHEM1, a hyperstable α-helical solenoid that employs an axial histidine ligand to coordinate iron protoporphyrin IX (FePPIX) with high affinity (Figure 1B)^32^. To replace the FePPIX with CoPPIX, we adapted a previously reported strategy for *in vivo* cofactor biogenesis and incorporation^35^ that obviates the need for costly *in vitro* cofactor replacement steps^36,37^. The UV-Vis spectrum of purified CoPPIX loaded dnHEM1 exhibits a diagnostic Soret feature at 422 nm, shifted from 402 nm as observed with FePPIX loaded dnHEM1, consistent with successful incorporation of CoPPIX^35^ (Supplementary Figure 1).

**Figure 1.**
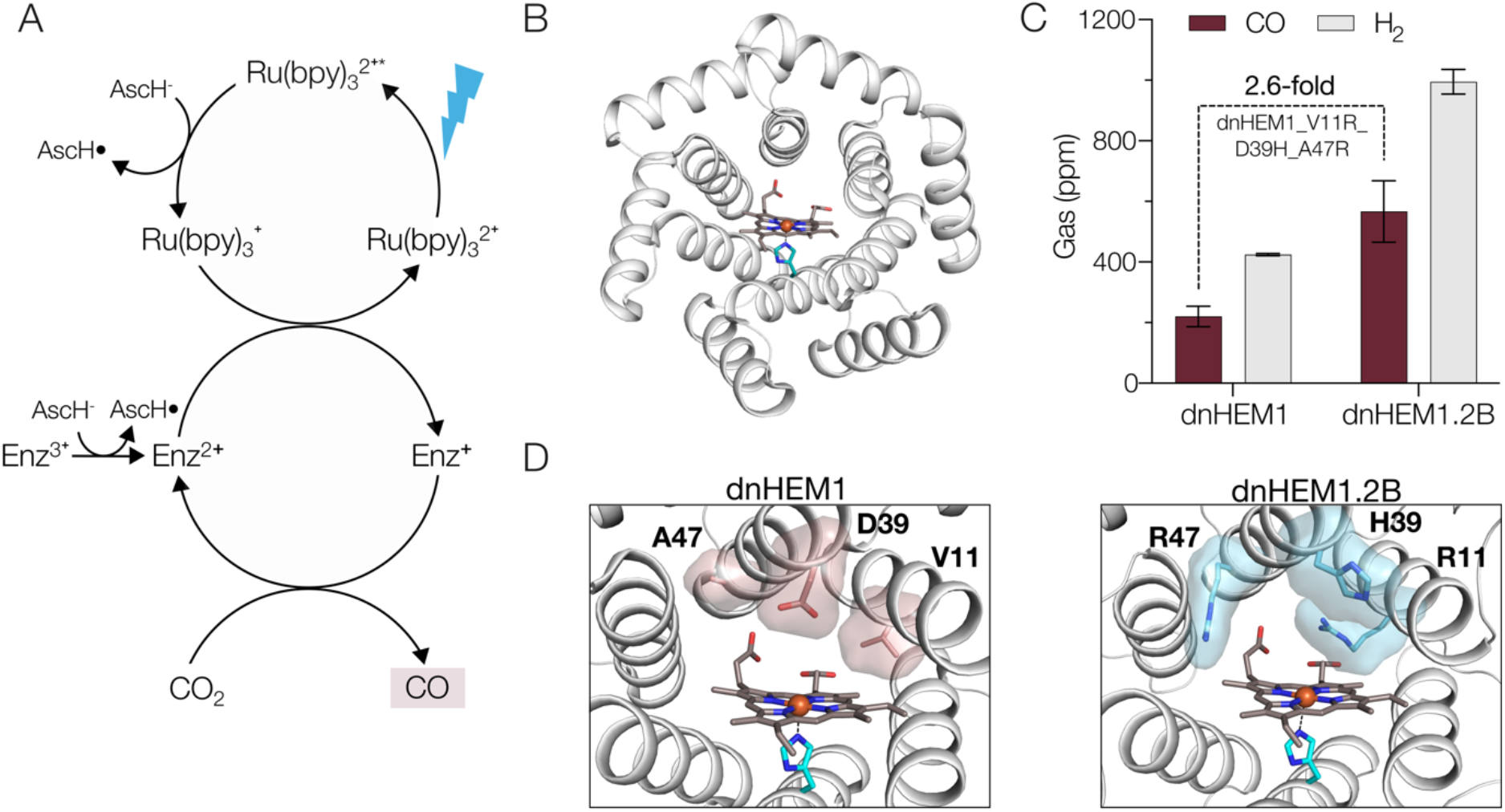
Development of first generation *de novo* CO_2_ reductases. A) Proposed photocatalytic cycle for enzymatic reduction of CO_2_ to CO using [Ru(bpy)_3_]^2+^ as a photosensitiser. AscH = sodium *L*-ascorbate. B) Crystal structure of dnHEM1 (PDB: 8C3W) highlighting the heme cofactor (atom-coloured sticks with dark grey carbons) and the axial histidine (atom-coloured sticks with blue teal carbons). C) Bar chart comparing the CO_2_ reductase activity of CoPPIX loaded dnHEM1 and dnHEM1.2B monitoring CO (plum) and H_2_ (grey) production. Biotransformations were performed using 0.3 μM holo-enzyme, 100 mM sodium L-ascorbate, 100 mM sodium bicarbonate, 100 µM [Ru(bpy)_3_]^2+^ in 500 mM potassium phosphate buffer (pH 7.0), and irradiation under blue light (475 nm) for 30 minutes. Error bars represent the range of values from measurements made in duplicate. D) Active sites of dnHEM1 (PDB: 8C3W) and dnHEM1.2B (design model) showing axial histidine (atom-coloured sticks with blue teal carbons) and heme cofactor (atom-coloured sticks with dark grey carbons), with the mutations between dnHEM1 and dnHEM1.2B highlighted in pink and blue, respectively.

We next evaluated the photocatalytic activity of dnHEM1 for CO_2_ reduction under anaerobic conditions, employing [Ru(bpy)_3_]^2+^ as a photosensitiser, sodium bicarbonate as a CO_2_ source and sodium *L*-ascorbate as a sacrificial reductant under blue light irradiation (475 nm) for 30 minutes (Figure 1A). Samples of the reaction headspace following photoirradiation were analysed by gas chromatography (GC). Under these conditions, dnHEM1 produces 220 ppm CO (94 turnovers in 30 min) alongside 424 ppm H_2_. In an attempt to further improve activity, we sampled an in-house panel of dnHEM1 variants that have previously been engineered to enhance heme peroxidase or carbene transferase activity^32^. From these assays, we identified dnHEM1.2B (dnHEM1_V11R_D39H_A47R, Figure 1D) that displays a 2.6-fold improvement in CO_2_ reductase activity (241 turnovers in 30 min), albeit with similar selectivity to that observed with dnHEM1 (Figure 1C).

### Design of enhanced CO_2_ reductases

Having developed a first generation *de novo* CO_2_ reductase, we next sought to explore a broader range of protein sequences and scaffolds to identify more potent and selective enzymes. We used multiple design strategies to generate heme binding proteins featuring either cysteine or histidine as an axial ligand (Figure 2A). Sequence diverse artificial homologs of myoglobin^38^ and dnHEM1.2B were generated using a combination of ProteinMPNN and RFjoint2 inpainting^39^. These approaches were extended to generate heme binders in *de novo* NTF2 scaffolds^40–42^. To explore a wider variety of protein topologies and active site configurations, we turned to generative AI based protein design methods. We have previously generated and characterized tens of *de novo* Cys-ligated heme-binding proteins based on RFdiffusion All-Atom protein backbone generation^33^. To expand this set, we started new design trajectories from a minimal active site model consisting of an axially ligated heme with a spatial placeholder in the distal site. The amino acid sequences were then optimized by iterative application of ProteinMPNN^43^, LigandMPNN^34^, Rosetta FastRelax^44^ and AlphaFold2^45^. Designed proteins were evaluated for AlphaFold2 confidence and accuracy, as well as for Rosetta metrics describing the protein-ligand interface quality. Synthetic DNA fragments encoding the designed proteins from each of the design pipelines were assembled into the pET29b plasmid for expression in *Escherichia coli*^46,47^. Following evaluation of the soluble expression levels and heme-binding properties of these designs, we selected a total of 69 well-performing sequences from 26 unique protein folds for CoPPIX incorporation and activity studies (Supplementary Table 1).

**Figure 2.**
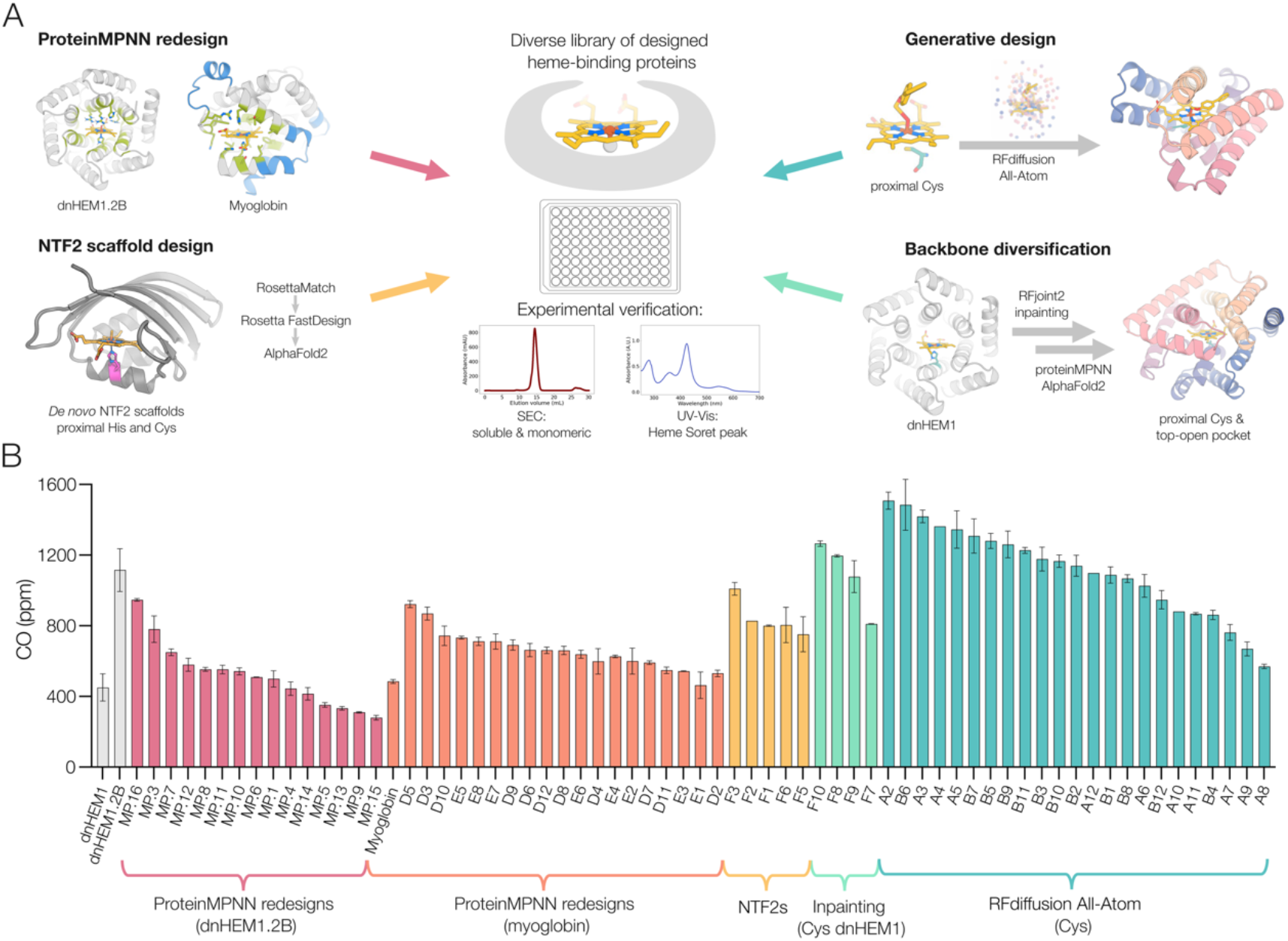
Design and screening of *de novo* CO_2_ reductases. A) Schematic of the heme-binding protein design pipelines using multiple *de novo* computational design methods. B) Bar chart displaying amount of CO (ppm) produced by the panel of CoPPIX loaded *de novo* designs compared to dnHEM1 and dnHEM1.2B (grey). The panel includes ProteinMPNN redesigns of dnHEM1.2B (pink), ProteinMPNN redesigns of myoglobin (orange), NTF2 scaffold-based designs (yellow), inpainting of Cys-ligated dnHEM1 designs (green), and designs generated by RFdiffusion All-Atom (teal). Biotransformations were performed using enzyme normalized to a λ_Soret_ absorbance of 0.1, 100 mM sodium *L*-ascorbate, 100 mM sodium bicarbonate, 100 µM [Ru(bpy)_3_]^2+^ in 500 mM potassium phosphate buffer (pH 7.0) and irradiated under blue light (475 nm) for 30 minutes at room temperature. Error bars represent the range of values from measurements made in duplicate.

The panel of designs was expressed in *E. coli* using the protocol described above for *in vivo* CoPPIX incorporation. Of the 69 sequences selected, 67 were expressed in soluble form and readily purified via nickel affinity chromatography. 63 of these designs displayed UV-Vis spectral features diagnostic of CoPPIX incorporation (λ_Soret_ values from 420 to 435 nm)^27,35,48– 50^ (Supplementary Figures 2-6). The remaining 4 designs have more complex UV-Vis absorbance profiles (Supplementary Figure 7), potentially rising from mixed oxidation states and/or partial spin states^35,51^.

The above 67 purified designs were subsequently evaluated for photocatalytic CO_2_ reduction activity using the headspace GC assay (Figure 2B). While sequence redesigns of dnHEM1.2B gave several variants with improved CO formation versus dnHEM1, this approach failed to yield any artificial homologs with improved activity compared to dnHEM1.2B. Similarly, redesigns based on myoglobin as a template (PDB: 3RGK) gave some improvements over the wild-type sequence but again failed to surpass the activity of dnHEM1.2B. Greater success was achieved using inpainting to introduce an axial cysteine ligand into dnHEM1 which gave small improvements over dnHEM1.2B. Evaluation of designs generated using RFdiffusion All-Atom incorporating a cysteine ligand led to the identification of eleven structurally diverse designs that displayed activity greater than or equal to dnHEM1.2B. Of the top performing designs, A2 and A3 share a common fold but had a propensity to precipitate during purification. We therefore selected A4 and B6, which display similarly high activity, for further characterisation. We note that these designs bear no resemblance to any protein observed in nature, as indicated by template modelling (TM) scores of 0.54 and 0.56, respectively^52^ (Supplementary Figure 8).

Following production of A4 and B6 under optimized expression conditions, the designs were purified by nickel affinity chromatography followed by size exclusion chromatography (SEC). SEC allowed separation of apo- and holo-proteins to enable isolation of fully occupied CoPPIX-loaded enzymes (Supplementary Figure 9). Both proteins eluted as monomeric species as confirmed by SEC-MALS (Supplementary Figure 10). UV-Vis spectroscopy showed that A4 exhibits a sharp Soret band at 434 nm with Q-band features at 542 and 577 nm, whereas B6 displays a Soret at 422 nm, with Q-band features at 533 and 563 nm (Figure 3A). CoPPIX incorporation was quantified using inductively coupled plasma optical emission spectrometry (ICP-OES), and used to determine Soret extinction coefficients of 104.4 and 97.4 mM^−1^cm^−1^ for A4 and B6, respectively (Supplementary Table 2). No iron contamination was detected in any sample by ICP-OES. A4 and B6 are both hyper-thermostable (Tm > 95 °C), showing minimal changes in circular dichroism (CD) spectra from 25–95 °C. Cofactor retention was also preserved at 95 °C, with negligible shifts in Soret intensity or wavelength (Supplementary Figure 11).

**Figure 3.**
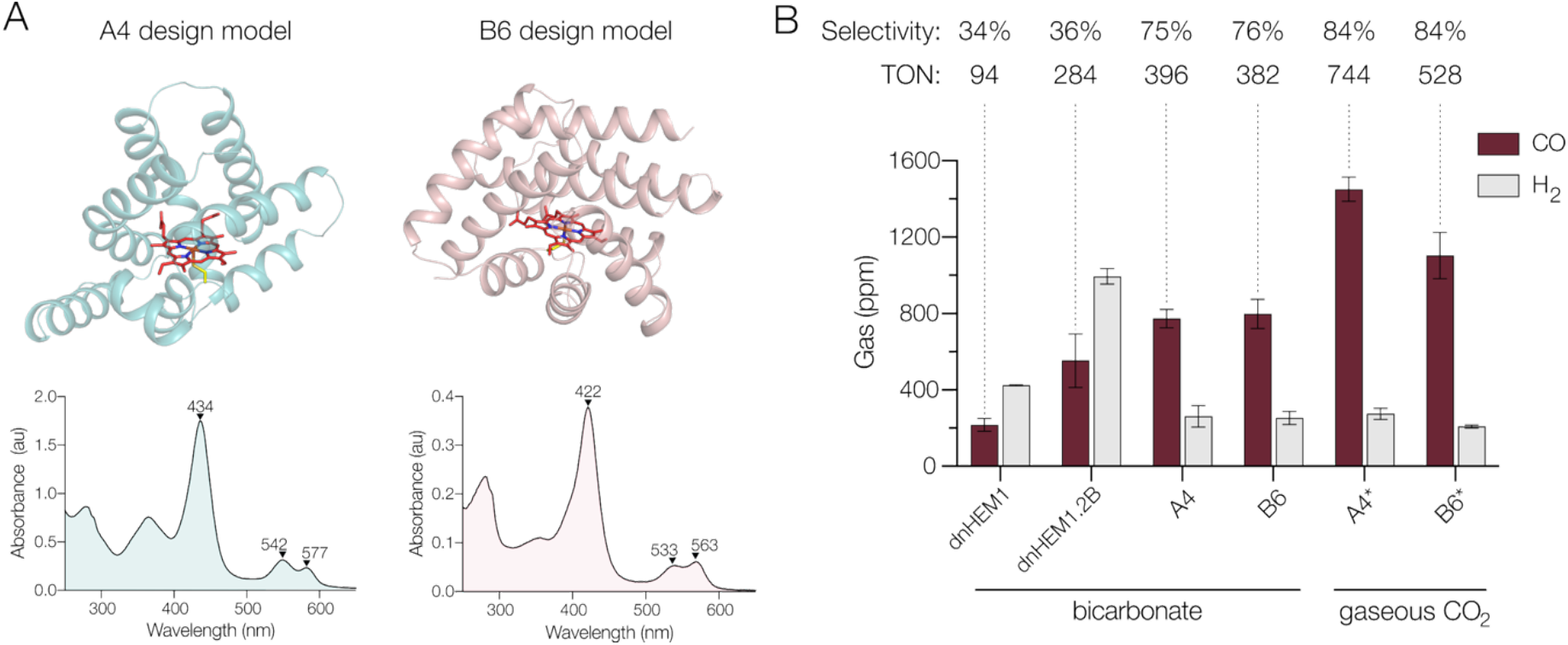
Characterisation of designs A4 and B6. A) AlphaFold2 predicted models of A4 (teal) and B6 (pink) showing the axially-ligated heme as red atom-coloured sticks. UV-Vis spectra of A4 (teal) and B6 (pink). B) Bar chart comparing the CO_2_ reductase (plum) and hydrogen evolution (grey) activity of dnHEM1, dnHEM1.2B, A4 and B6. Biotransformations were performed using 0.3 μM holo-enzyme, 100 mM sodium *L*-ascorbate, 100 µM [Ru(bpy)_3_]^2+^ in 500 mM potassium phosphate buffer (pH 7.0) with either 100 mM sodium bicarbonate or gaseous CO_2_ as the CO_2_ source, and irradiation under blue light (475 nm) for 30 minutes. Error bars represent the range of values from measurements made in duplicate.

With purified enzymes in hand, we next compared the activity and selectivity of A4, B6, dnHEM1, and dnHEM1.2B. Consistent with the initial panel screening, A4 and B6 exhibited 3.6- and 3.7-fold improvements in CO_2_ reduction activity compared to dnHEM1, respectively. These activity gains were accompanied by a marked increase in selectivity for CO_2_ reduction over H_2_ generation (*ca*. 75 % selectivity with A4 and B6 *vs. ca*. 35 % with dnHEM1 and dnHEM1.2B) (Figure 3B). It is known that photoactivated [Ru(bpy)_3_]^2+^ can drive H_2_ evolution^53^, and control experiments show that hydrogen production is significantly reduced in the presence of A4 and B6 (Supplementary Figure 12). These observations suggest that the intrinsic selectivity associated with A4 and B6 is likely substantially higher than 75%. Time-course experiments show that A4 and B6 can achieve TTN_*CO*_ values of 580 and 620, prior to deactivation (Supplementary Figure 13). Improvements in reaction rate, selectivity, and total turnovers can be achieved by replacing bicarbonate with gaseous CO_2_ as the substrate. Using A4 and B6 as biocatalysts, CO_2_ can be converted to CO with >84% selectivity, at a rate of 25 min^-1^ and 18 min^-1^, respectively, and >1000 total turnovers was achieved for each prior to enzyme deactivation (Figure 3B, Supplementary Table 3, and Supplementary Figure 13).

### Structural analysis

We next sought to solve structures of A4 and B6 for comparison to the design models. While B6 proved difficult to crystallize, the crystal structure of A4 was determined to a resolution of 2.1 Å (PDB: 9T8A). This structure closely agrees with the design model and with the AlphaFold2 model of A4, with a backbone RMSD of 1.05 Å (Figure 4A). The CoPPIX cofactor resides in a solvent-occluded pocket (solvent accessible surface area of 12 Å^2^ *vs*. 112 Å^2^ for dnHEM1, Supplementary Figure 14) and is ligated through Cys98 as intended. However His73, designed to interact with one of the CoPPIX propionate groups, instead occupies the vacant coordination site to generate a hexa-coordinated state, which also results in a minor displacement of the porphyrin cofactor relative to the model. The His73 conformer observed in the x-ray structure was occluded in the design model by the surrogate distal site ligand used as a placeholder in the design pipeline (Figure 4B). In the A4 crystal structure, the porphyrin also adopts a rotated geometry relative to the design model, leading to polar interactions between the CoPPIX propionate and Thr64. An AlphaFold3^54^ model of A4 with heme agrees with the crystal structure both regarding the orientation of the porphyrin, as well as the His73 conformer (Supplementary Figure 15).

**Figure 4.**
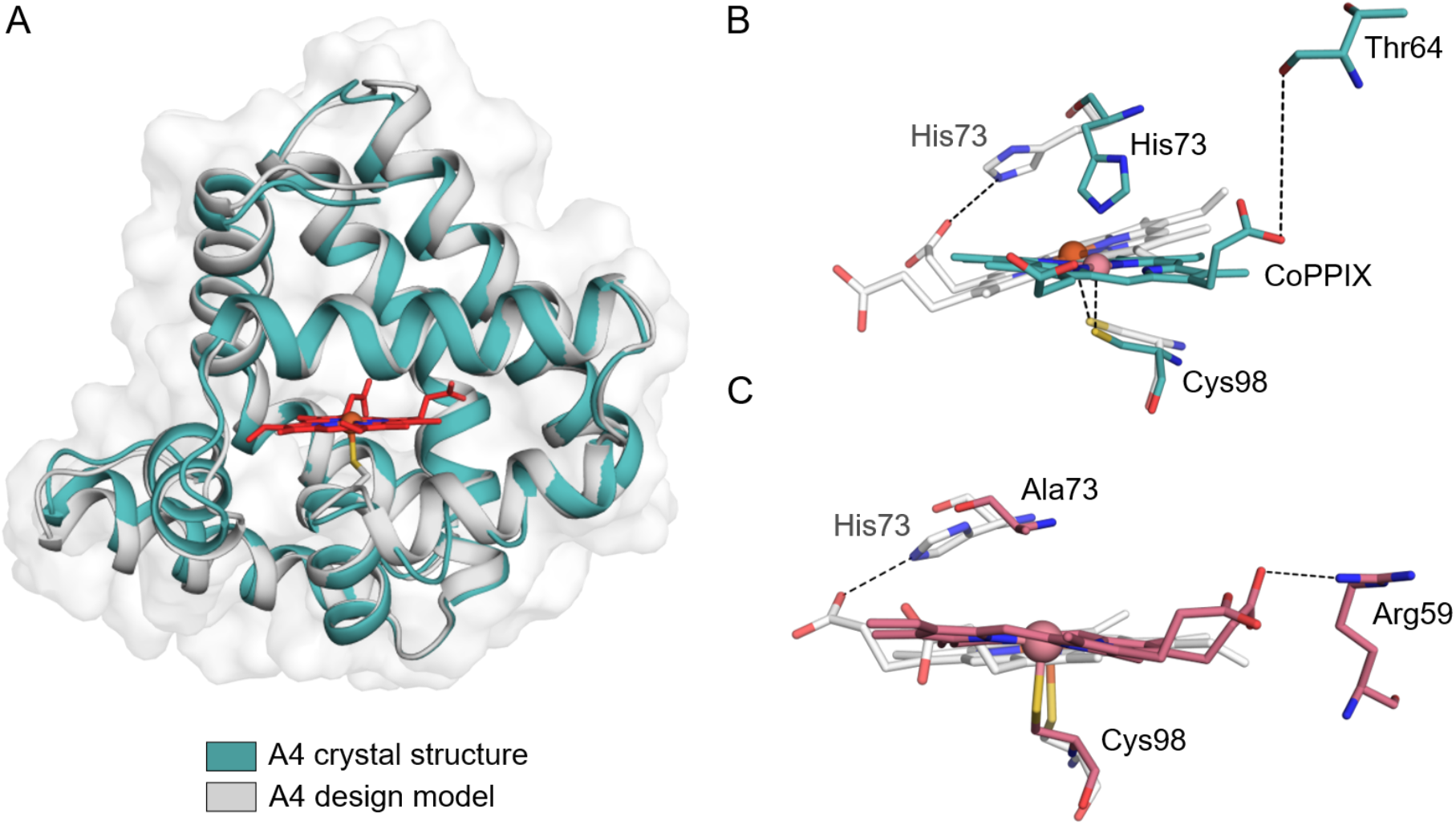
Structural Analysis of A4 and A4 H73A. A) A crystal structure of A4 (teal) (PDB: 9T8A) shows close agreement to the AlphaFold2 design model (grey). The cysteine-ligated heme of the design model is shown as red atom-coloured sticks. B) Comparison of the active site of A4 (teal atom-coloured sticks) and the A4 design model (grey atom-coloured sticks) highlighting the differences in positioning of His73 and the porphyrin cofactor. In the design model, His73 forms interactions with the propionate group of the cofactor, whereas in the crystal structure it coordinates the CoPPIX cofactor to form a hexa-coordinated complex. C) Comparison of the active site of A4 H73A (PDB: 9T8B) (pink atom-coloured sticks) and the A4 design model (grey atom-coloured sticks). In the A4 H73A structure the CoPPIX cofactor is penta-coordinated as required for catalysis, with the porphyrin plane and positioning of the axial cysteine now more closely resembling the design model.

To understand the functional significance of Cys98 and His73, we independently mutated these residues to alanine. Substitution of Cys98 led to a decrease in CoPPIX binding, activity and selectivity, demonstrating that the programmed cysteine ligand plays a key role in CoPPIX-dependent CO_2_ reduction (Supplementary Figures 16 & 17). In contrast, the impact of the His73A mutation was found to be negligible. A 1.8 Å resolution structure of A4 H73A (PDB: 9T8B) shows that while the porphyrin still adopts the rotated geometry observed in A4, its plane and the positioning of the axial Cys98 ligand now more closely resemble that of the design model. In this structure, the CoPPIX cofactor is now penta-coordinated as required for catalysis (Figure 4C).

### Conclusions

In summary, this study illustrates how deep learning-based protein design can be used to generate artificial metalloenzymes to transform difficult-to-activate molecules such as CO_2_. By sampling designs with diverse topologies, active-site architectures and cofactor coordination environments, we identified several enzymes featuring a Cys-ligated CoPPIX cofactor that display enhanced CO_2_ reductase activity while minimizing competing off-pathway H_2_ evolution processes. With limited guiding principles to steer active site design, and technically challenging experimental protocols that preclude standard directed evolution methods to optimize enzyme activity, these discoveries would not have been possible without access to a diverse set of protein scaffolds hosting the catalytic cofactor. Moving forward, there are opportunities to extend our enzyme design pipeline to achieve more valuable conversions of CO_2_ into C_2_ and C_3_ platform chemicals that may also be more amenable to high-throughput quantification to underpin directed evolution workflows. In this way, we aim to deliver a new generation of efficient, selective and stable CO_2_-transforming enzymes to help achieve our net-zero goals.

## Acknowledgements

We would like to acknowledge funding from EPSRC and bp through the bp International Centre for Advanced Materials (bp-ICAM) for an ICASE to E.R. (EP/W522065/1), a Prosperity Partnership funded by EPSRC, bp through the bp-ICAM (grant numbers ICAM75, ICAM80 and ICAM83) and Johnson Matthey plc to E.R. and A.A. (EP/V056565/1), and the Engineering and Physical Sciences Research Council to A.P.G. and D.B. (EPSRC Centre-to-Centre Partnership, EP/Z531157/1). We are grateful to the Manchester SYNBIOCHEM Centre (BB/M017702/1), the Future Biomanufacturing Hub (EP/S01778X/1) and the Henry Royce Institute for Advanced Materials (funded through EPSRC grants nos. EP/R00661X/1, EP/S019367/1, EP/P025021/1 and EP/P025498/1) for access to their facilities. We acknowledge assistance given by Research IT and the use of the Computational Shared Facility at the University of Manchester. We thank J. Tibble-Howlings from the Analytical Services and Instrumentation at the University of Manchester for their expertise and support in ICP-OES measurements. The authors would like to thank Diamond Light Source for beamtime on i03 and i04 (proposal MX38021-12 & MX31850-102). We thank the Open Philanthropy Project Improving Protein Design Fund (D.B., I.K.), the Howard Hughes Medical Institute (D.B., I.K.), MCIN/AEI/10.13039/501100011033 (grants PID2024-160774OB-I00, PID2021-125946OB-I00; R.N.F), and NERSC for access to the Perlmutter high-performance computing resources (award BER-ERCAP0022018). We thank Luki Goldschmidt and Kandise VanWormer for maintaining the computational and wet-lab resources at the Institute for Protein Design. We thank Xinting Li for assistance with mass spectrometry experiments at the Institute for Protein Design. We thank Dr. Florence Hardy and Dr. Nathan Ennist for helpful discussions, Dr. Christopher Norn for providing the *de novo* designed NTF2 scaffolds, and Dr. David Juergens for assistance with RFjoint2 inpainting.

## Methods

### Design of a library of heme-binding proteins

The library of diverse heme-binding proteins was constructed using several different design methods, with the common goal of creating proteins with a proximal histidine or cysteine ligation, and an open distal pocket. The following design strategies were used: (1) Creating diverse sequence analogs of dnHEM1.2B peroxidase with ProteinMPNN. The design of these proteins is described in more detail in Supplementary Methods, section 3.2. (2) Installing heme binding sites into previously designed nuclear transport factor 2 (NTF2) protein scaffolds using Rosetta-based sequence design^40,55,56^ and AlphaFold2 structure prediction. The design and characterization of these proteins are described in more detail in Supplementary Methods, section 3.3. (3) Creating diverse sequence and structural analogs of myoglobin^57^ (PDB: 3RGK) with ProteinMPNN and RFjoint2 inpainting. The design and characterization of these variants have been previously reported^38^. (4) Structurally modifying a *de novo* designed heme-binder, dnHEM1, with RFjoint2 inpainting to incorporate a proximal Cys residue and to create alternative pocket opening geometries. The design and characterization of these proteins are described in more detail in Supplementary Methods, section 3.4. (5) Creating new protein scaffolds completely from scratch using RFdiffusion All-Atom. These proteins were created starting from a heme model with a proximal cysteine residue and a distal substrate placeholder. ProteinMPNN, LigandMPNN, Rosetta FastRelax and AlphaFold2 were used to design and validate the sequences corresponding to the generated protein backbones. A subset of these designs have previously been reported^33^. Additional newly characterized variants are described in Supplementary Methods, section 3.5. Supplementary Table 1 lists the plate layout of the heme-binder library. Amino acid and DNA sequences are displayed in Supplementary Table 4.

### Materials

All materials were obtained from commercial suppliers and used as received, unless otherwise stated. Chemicals were sourced from Sigma-Aldrich unless otherwise indicated. LB agar, LB media, 2×YT media, isopropyl-β-*d*-1-thiogalactopyranoside (IPTG) and *L*-arabinose were purchased from Formedium. Terrific Broth II (TB-II, MP Biomedicals) powder was purchased from Fisher Scientific. *Escherichia coli* strains BL21 (DE3) and 5α, Q5 DNA polymerase, T4 DNA ligase and restriction enzymes were purchased from New England Biolabs. *Escherichia coli* strain BL21-AI and sodium bicarbonate were purchased from Thermo Fisher. Oligonucleotides and genes were synthesized by Integrated DNA Technologies (IDT).

### Construction of variants

Genes encoding the design panel were purchased from IDT and assembled (Golden Gate cloning) into pET29b(+) vector (LM627, AddGene 191551)^46^ with a C-terminal SNAC cleavage site^58^ and subsequent hexa-histidine tag. The final expressed amino acid sequences were obtained as MSG-design-GSGSHHWGSTHHHHHH. Constructs without a SNAC-cleavage site were generated using overlap extension PCR and subcloned into a pET29b(+) vector. Variants (A4 H73A and A4 C98A) were created using the QuikChange Lightning Mutagenesis kit (Agilent) according to the published protocol. All constructs were verified using Sanger sequencing (Eurofins).

### Protein expression and purification

Chemically competent *E*.*coli* BL21-AI cells were transformed with the appropriate plasmid encoding the designed protein. Single colonies of freshly transformed cells were cultured for 18 h in 10 mL modified M9 minimal media to minimize heme contamination (1x M9 salts, 0.2% glucose, 0.1 mM CaCl_2_, 2 mM MgSO_4_, 4 mg mL^-1^ casamino acids, 10 μg mL^-1^ thiamine chloride), supplemented with 25 μg ml^-1^ kanamycin. Starter cultures were used to inoculate 400 mL modified M9 media supplemented with 25 µg mL^−1^ kanamycin. Expression of *in vivo* CoPPIX-loaded proteins was adapted from a previously reported strategy^35^. Cells were grown at 37 °C with shaking at 180 rpm until OD_600_ of 0.8. At this point, 1 mM δ-aminolevulinic acid (δ-ALA) and 500 mM CoCl_2_ were added. After an additional 30 minutes, temperature was reduced to 25 °C, and gene expression was induced with 1 mM IPTG and 0.2% *L*-arabinose, and grown for a further 18 hours. Cells were harvested by centrifugation (6880 x *g*, 4 °C for 10 min).

Pelleted cells were resuspended in lysis buffer (50 mM KPi, 200 mM NaCl, 20 mM imidazole, pH 7.5) supplemented with 1 mg mL^-1^ lysozyme and 1 mg mL^-1^ DNase. Cells were lysed by sonication (13 mm probe, 1s on, 1s off, 50% amplitude, total duration 20 mins). The lysate was clarified by centrifugation at 10,000 × *g* for 30 mins. The supernatant was incubated with nickel-nitrilotriacetic acid (Ni-NTA) resin for 1 hour at 4 °C, then packed onto a gravity flow column. Proteins were eluted using a high imidazole buffer (250 mM imidazole, 50 mM KPi, 200 mM NaCl, pH 7.5). Proteins were concentrated by centrifugation using Vivaspin® 20, 10 kDa MWCO. Subsequently, designs were further purified by SEC ÄKTApure system using a Superdex 75 16/600 column (GE Healthcare) pre-equilibrated with buffer (50 mM KPi, 200 mM NaCl pH 7.0, Supplementary Figure 9). Proteins were aliquoted, flash frozen in liquid nitrogen and stored at -80 °C.

### Spectrophotometric assays

Analysis of the UV-Visible (UV-Vis) spectroscopic properties were carried out using an Agilent Cary 8485 UV-Vis spectrophotometer (CoPPIX-loaded proteins), or Jasco Spec V-750 UV-Vis spectrophotometer and BioTek Synergy Neo2 microplate reader (heme-loaded proteins).

### Size Exclusion Chromatography Multi-Angle Light Scattering (SEC-MALS)

The oligomeric state of A4 and B6 were assessed using SEC-MALS. Purified samples (0.2 mg mL^-1^) were loaded onto a Superdex 75 (GE Healthcare) at 0.75 mL min^-1^ with a constant flow of storage buffer (50 mM KPi, 200 mM NaCl, pH 7.0). The samples were passed through a Wyatt Heleos II EOS 18-angle laser photometer coupled to a Wyatt Optilab rEX refractive index detector. Data analysis was performed (Hydrodynamic radii and molecular mass measurements were analysed) using ASTRA 6 software (Wyatt Technologies).

### Circular dichroism

To determine the thermostability of the designs, far-ultraviolet circular dichroism (CD) measurements were carried out on an Applied Photophysics Chiroscan CD instrument. The 200 to 260 nm wavelength scans were measured at 10 °C intervals from 25 °C to 95 °C, using 0.3 mg mL^-1^ enzyme in 50 mM KPi (pH 7.0) in a 1 mm path length cuvette. Spectra were acquired once the temperature had stabilized to within 0.1 °C of the target temperature for 30 seconds. The same conditions were also employed to collect UV-Vis spectra between 250 and 700 nm for protein samples with an absorbance of ∼1.0 in the Soret region in a 1 cm path length cuvette.

### Determination of extinction coefficients

The extinction co-efficient of selected variants were determined using a combination of UV-Vis spectroscopy and ICP-OES as previously reported^28,59^. The Soret absorbance of variants were measured on an Agilent Cary 8485 UV-Vis spectrophotometer. The concentration of cobalt within the samples were measured using an AnalytikJena PlasmaQuant PQ 9000 Elite High-Resolution ARRAY ICP-OES (argon axial mode and plasma power at 1200W). Samples were prepared by adding highly concentrated protein stocks (50-200 µL) to HPLC-analysis grade nitric acid (1 mL) and stirred at 70 °C overnight. Samples were diluted with HPLC ultrapure grade water to a final volume of 10 mL (5% nitric acid final concentration) and passed through a 0.22 μm filtration system. Cobalt standard solutions were prepared from a 1000 ppm atomic absorption standard in 5% nitric acid (calibration curve from 0.625 to 10 ppm). Cobalt was measured at 237.863 nm and metal content was quantified by comparison to a standard curve (Supplementary Table 2).

### Mass spectrometry

Intact protein masses were acquired on a 1200 series LC QTOF 16600 MS (Agilent). Protein samples (1 mg mL^-1^) were injected on the LC-MS system, and desalted using a PLRP-S cartridge and eluted over a 6 minute 5% - 95% ACN gradient at 0.8 mL min^-1^. The resulting multiply charged spectrum was deconvoluted using the Agilent MassHunter BioConfirm software. Data presented in (Supplementary Table 5).

### Photoreduction CO_2_ assays

We evaluated a series of designed heme binding proteins for photocatalytic CO_2_ reduction activity. The panel comprises designed heme binding proteins previously reported^33,38^, alongside a selection of new designs (A3-A7 and B6-B7) that have not been disclosed previously. All reactions contained enzyme at the stated concentrations, 100 mM sodium *L*-ascorbate, 100 mM sodium bicarbonate (as CO_2_ source), 100 µM [Ru(bpy)_3_]^2+^ in 500 mM potassium phosphate buffer at pH 7.0, to a final volume of 2 mL, within 10 mL gas tight vials under anaerobic conditions. When pure CO_2_ gas was used, the vials were purged for 60 seconds each. Reactions were initiated by irradiation at 475 nm with blue LED light for the amount of time stated.

Samples were then removed from the light and left to equilibrate in the dark for 30 minutes. Directly after, 2 mL of the head space sample was injected onto an Agilent 490 Micro Gas Chromatography (GC) equipped with a Thermal Conductivity Detector (TCD) and a MS15A column (Supplementary Figure 18). The inlet injector temperature was set at 110 °C, column temperature at 80 °C and column pressure at 150 kPa. The detector response to CO and H_2_ was quantified using calibration curves made using gas analytic standards containing fixed amounts of CO, H_2_, and Ar (Supplementary Figure 19).

The number of moles of CO (or H_2_) in the standard gas injections was calculated by first rearranging the ideal gas equation. Turnover number (TON) was calculated by dividing the moles of CO or H_2_ gas detected by the moles of protein. Selectivity for CO production was calculated according to the following equation:

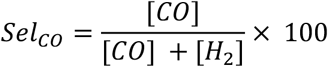

### Protein crystallization and refinement

Crystals of A4 and A4 H73A were obtained from a pET29b(+) construct lacking the SNAC-tag but retaining the C-terminal His-tag. SEC purified A4 and A4 H73A were concentrated to 7 mg mL^-1^ in 50 mM HEPES buffer (pH 7.5). Crystallization conditions for A4 and A4 H73A were identified using the BCS screen (Molecular Dimensions) and LMB screen (Molecular Dimensions), respectively.

Crystals suitable for diffraction experiments were obtained by sitting drop vapor diffusion at 4 °C in 400 nL drops containing equal volumes of protein and precipitant using a nano-dispenser (Mosquito TTP Labtech). Crystals of A4 was found in BCS A8: 0.1 M Phosphate/Citrate pH 5.5 20 % v/v PEG Smear High. Crystals of A4 H73A were found in LMB C9: 26 % w/v PEG 2000 MME, 0.1 M Bis-Tris pH 5.8. Before data collection, crystals were cryoprotected by the addition of 20% PEG 200 to the mother liquor and plunge-cooled in liquid nitrogen.

All data were collected on beamline iO3 (wavelength 0.9763 Å) at the Diamond Light Source Facility (Harwell, UK) and reduced and scaled with Xia2 DIALS. Structures were solved by molecular replacement using the AlphaFold2^45^ predicted model as the search model. Iterative rounds of model building and refinement were performed in Coot^60^ and phenix.refine^61^. Validation with MOLPROBITY^62^ and PDBREDO^63^ were incorporated into the iterative rebuild and refinement process. Data refinement statistics are shown in Supplementary Table 6. The coordinates and structure factors have been deposited in the Protein Data Bank under accession numbers 9T8A and 9T8B.

## Data Availability

The data generated in this study are provided within the paper and in the Supplementary Information. Source Data are provided with this paper. The coordinates and structure factors for the crystallographic data in this study are available in the Protein Data Bank under accession numbers 9T8A and 9T8B for A4 and A4 H73A respectively.

