## Supplementary Information for "A *de novo* CO_2_ Reductase Featuring a Cysteine-Ligated Cobalt Porphyrin Cofactor"

[a] Department of Chemistry, University of Manchester, Manchester Institute of Biotechnology, Manchester, UK.
[b] Institute for Protein Design, University of Washington, Seattle, USA; Department of Biochemistry and Howard Hughes Medical Institute, University of Washington, Seattle, USA.
[c] Department of Chemistry, University of Texas at Austin, Austin, USA.
† These authors contributed equally.

### 1. Supplementary Figures

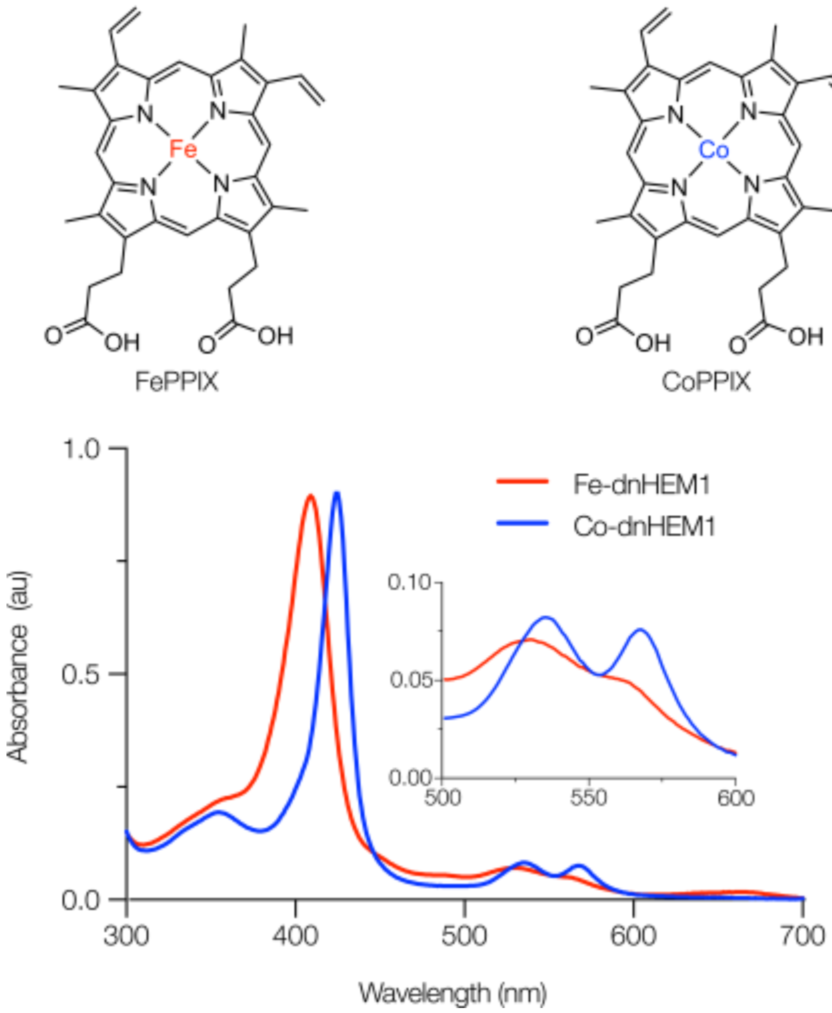

**Supplementary Figure 1**. CoPPIX loading in dnHEM1. Above: chemical structures of FePPIX and CoPPIX. Below: UV-Vis spectra of dnHEM1 loaded with FePPIX *in vitro* with Soret feature at 402 nm (Fe-dnHEM1, red) and dnHEM1 loaded with CoPPIX *in vivo* with Soret feature at 422 nm (Co-dnHEM1, blue). Insert shows Q-band region of Fe-dnHEM1 (528 and 563 nm) and Co-dnHEM1 (534 and 567 nm).

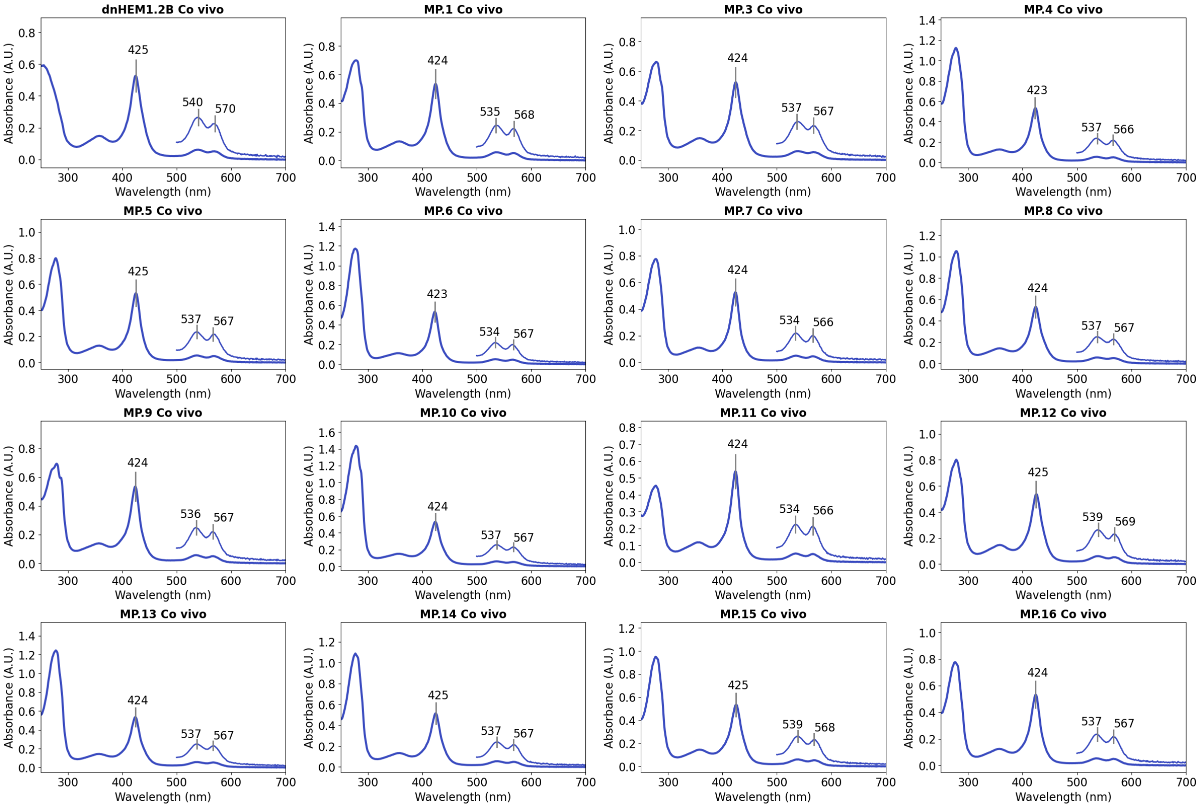

**Supplementary Figure 2.** UV-Vis spectra of *in vivo* CoPPIX-loaded ProteinMPNN-redesigns of dnHEM1.2B. Inset shows the visible region at 4x magnification. Spectra were recorded in a buffer containing 50 mM potassium phosphate, 200 mM NaCl at pH 7.2.

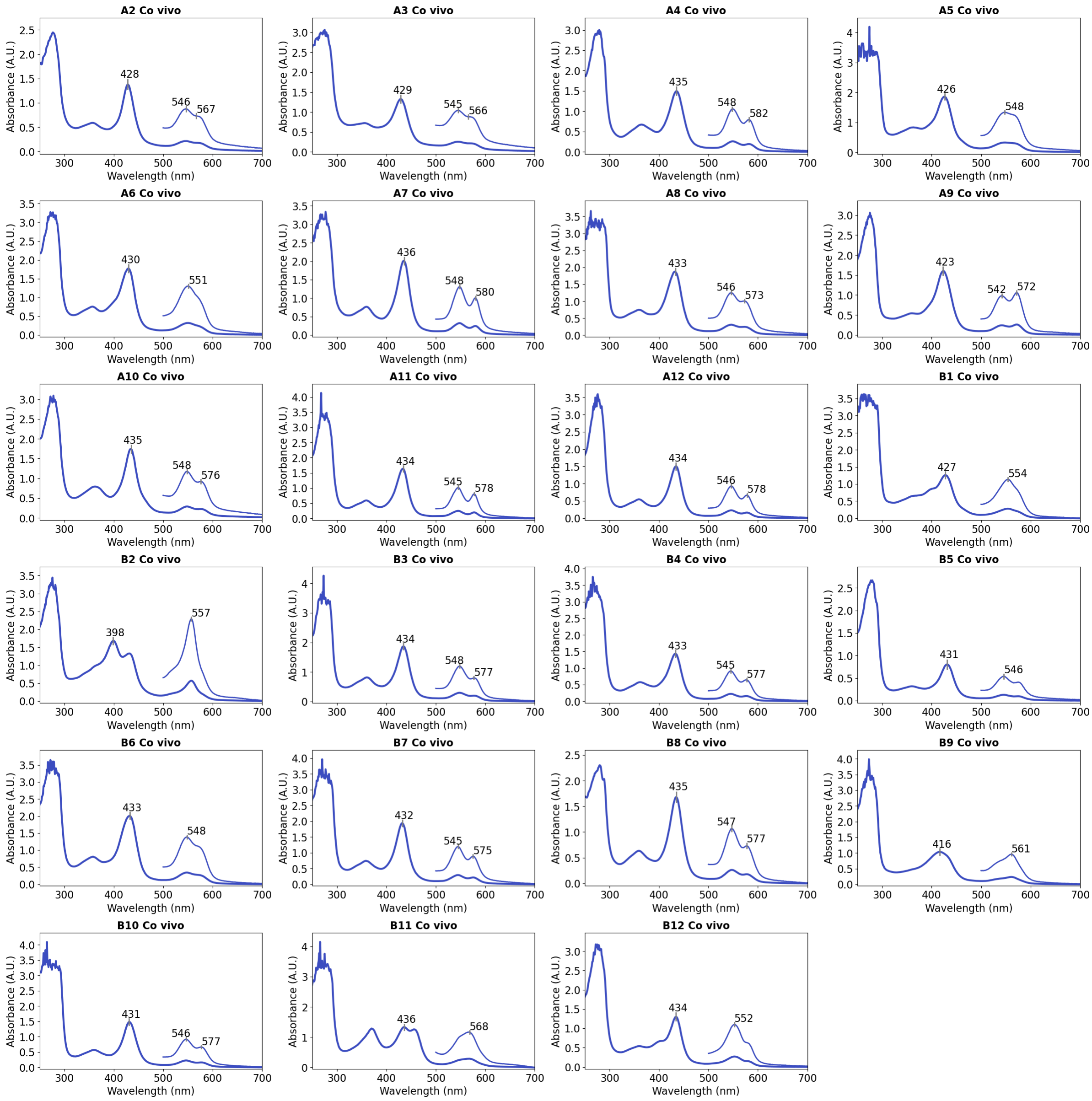

**Supplementary Figure 3**. UV-Vis spectra of *in vivo* CoPPIX-loaded proteins designed with RFdiffusion All-Atom. Inset shows the visible region at 4x magnification. Spectra were recorded in a buffer containing 50 mM potassium phosphate, 200 mM NaCl at pH 7.2.

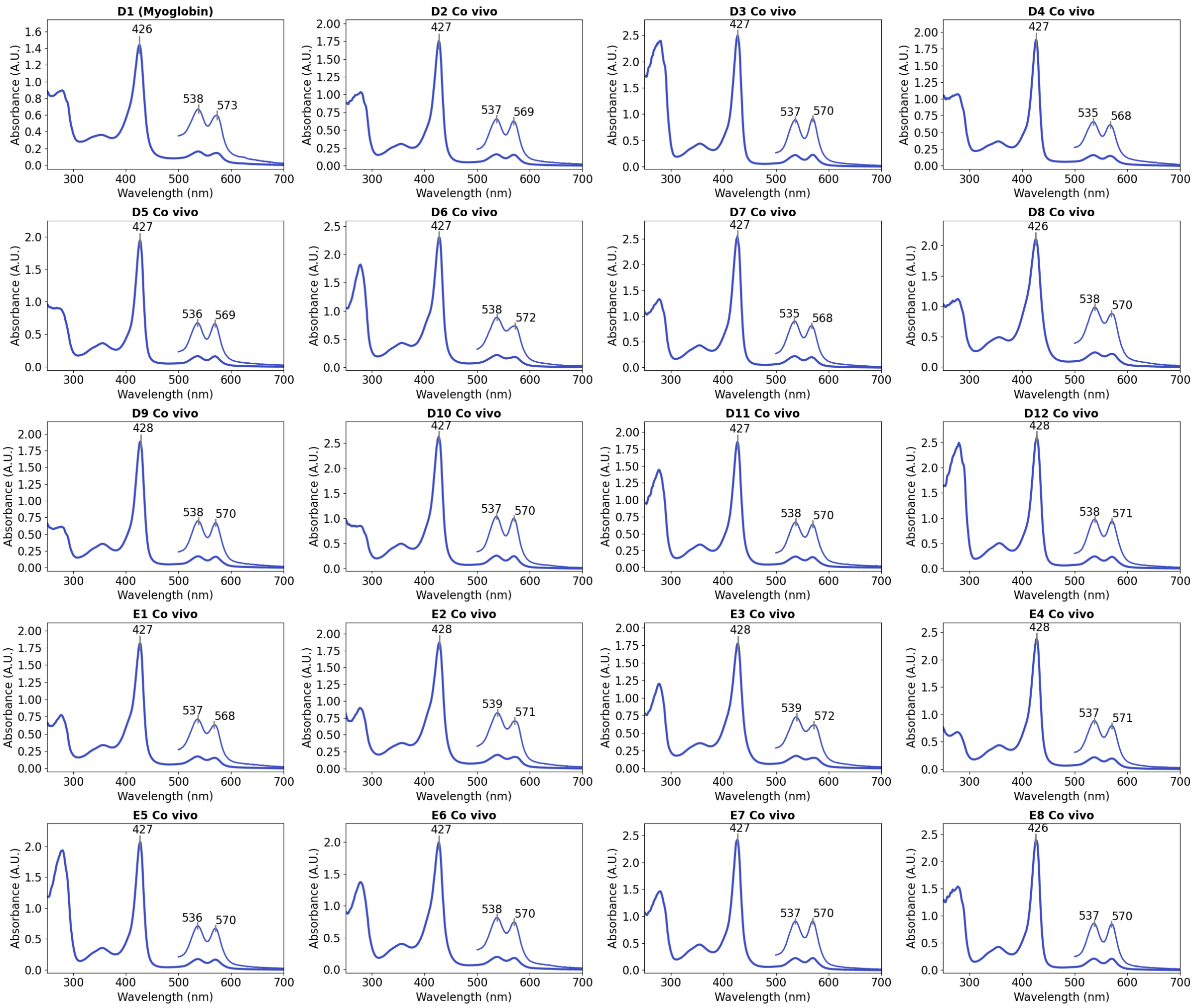

**Supplementary Figure 4.** Supplementary Figure 4. UV-Vis spectra of *in vivo* CoPPIX-loaded redesigned myoglobin variants. Inset shows the visible region at 4x magnification. Spectra were recorded in a buffer containing 50 mM potassium phosphate, 200 mM NaCl at pH 7.2.

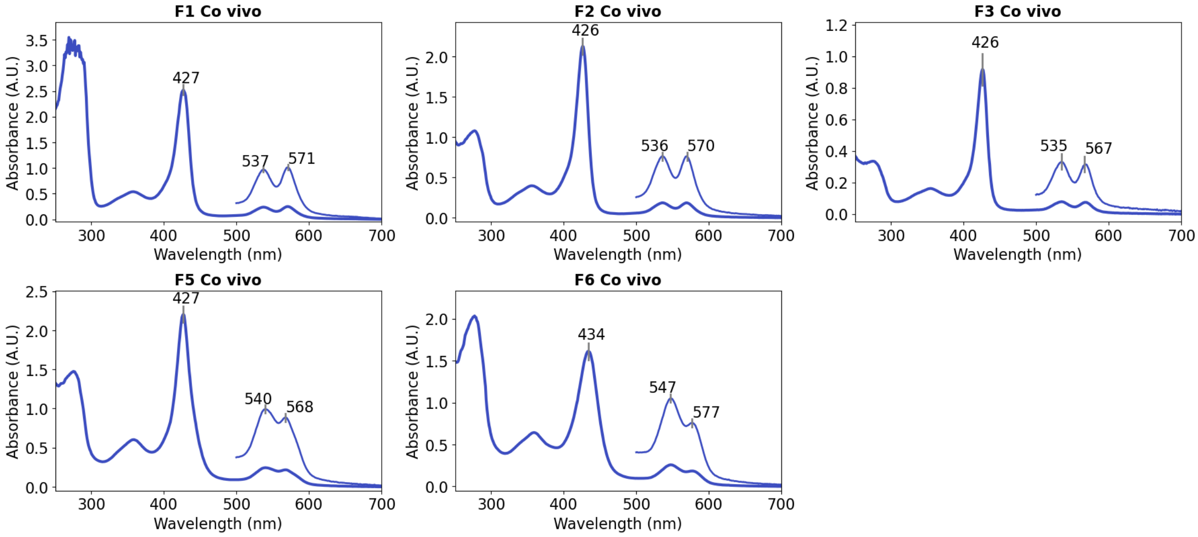

**Supplementary Figure 5**. UV-Vis spectra of CoPPIX-loaded NTF2HEM proteins. Inset shows the visible region at 4x magnification. Spectra were recorded in a buffer containing 50 mM potassium phosphate, 200 mM NaCl at pH 7.2.

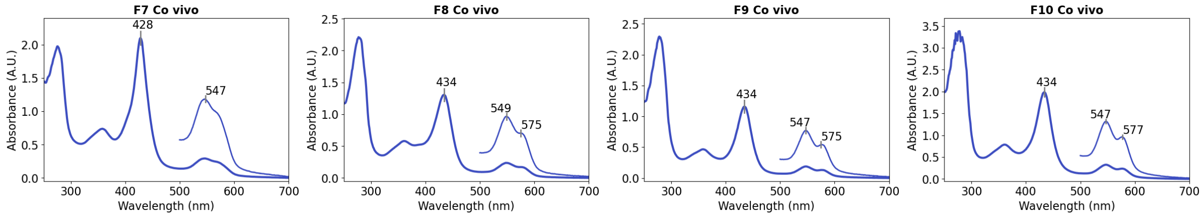

**Supplementary Figure 6**. UV-Vis spectra of CoPPIX-loaded inpainted Cys-dnHEM1 proteins. Inset shows the visible region at 4x magnification. Spectra were recorded in a buffer containing 50 mM potassium phosphate and 200 mM NaCl at pH 7.2.

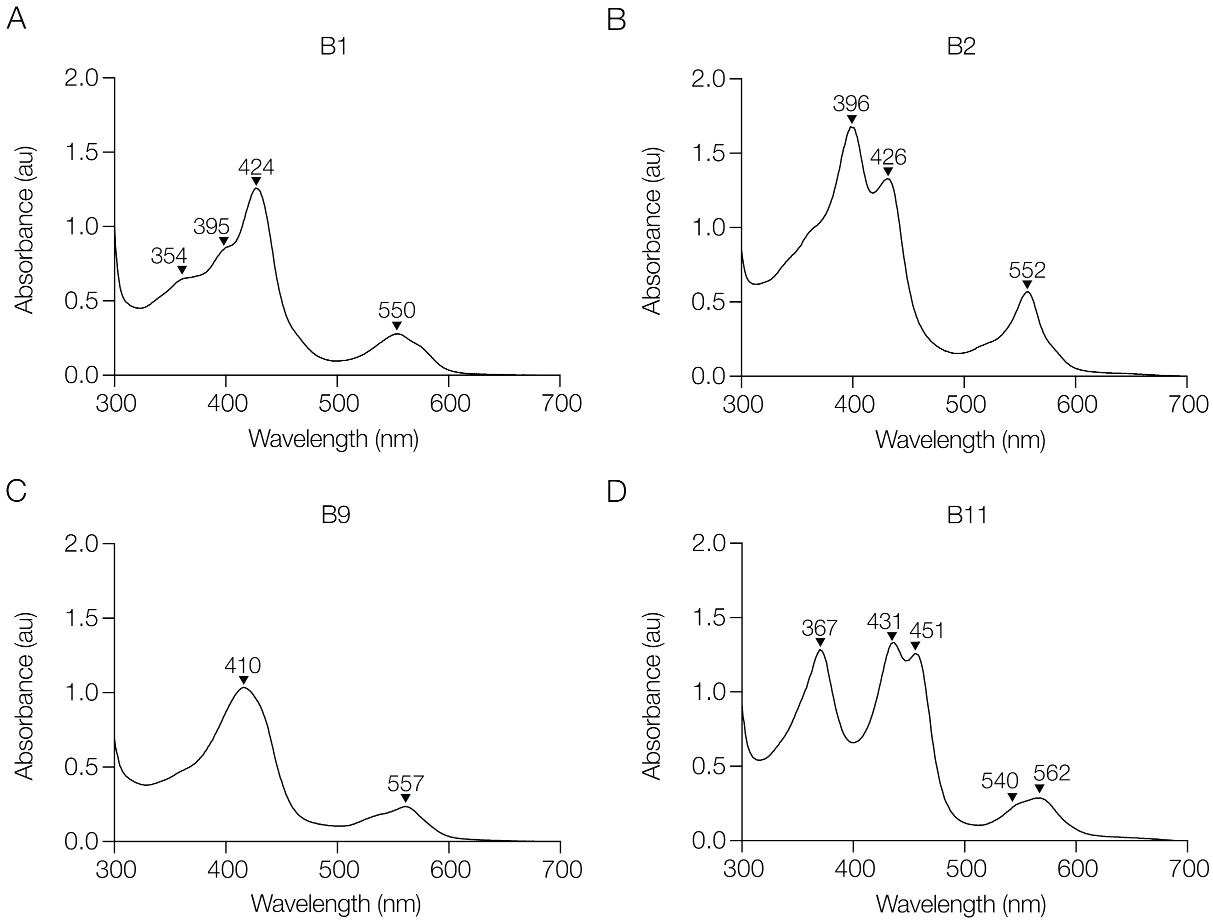

**Supplementary Figure 7.** Unusual spectra features of a subset of *in vivo* CoPPIX-loaded proteins designed by RFdiffusion All-Atom. A) B1. B) B4. C) B9. B) B11. Spectra were recorded in a buffer containing 50 mM potassium phosphate, 200 mM NaCl at pH 7.2.

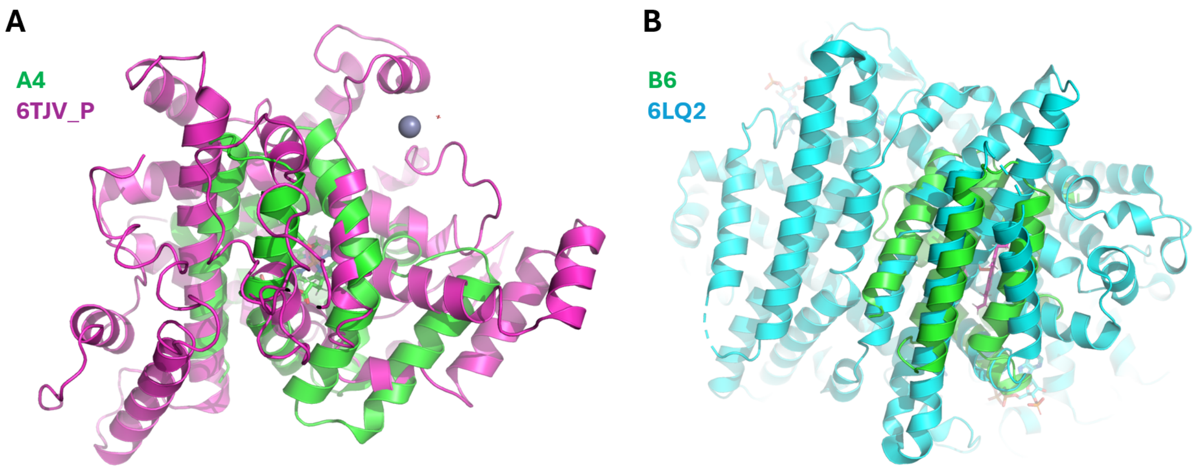

**Supplementary Figure 8.** Comparison of designs with closest matches from the PDB, based on the template modelling score calculated with TMalign[1]. Neither of the matches contains heme, and the alignments occur only through a few disconnected secondary structure elements. A) A4 aligned to PDB id 6TJV, chain P (TM score = 0.54). B) B6 aligned to PDB id 6LQ2, chain A (TM score = 0.56).

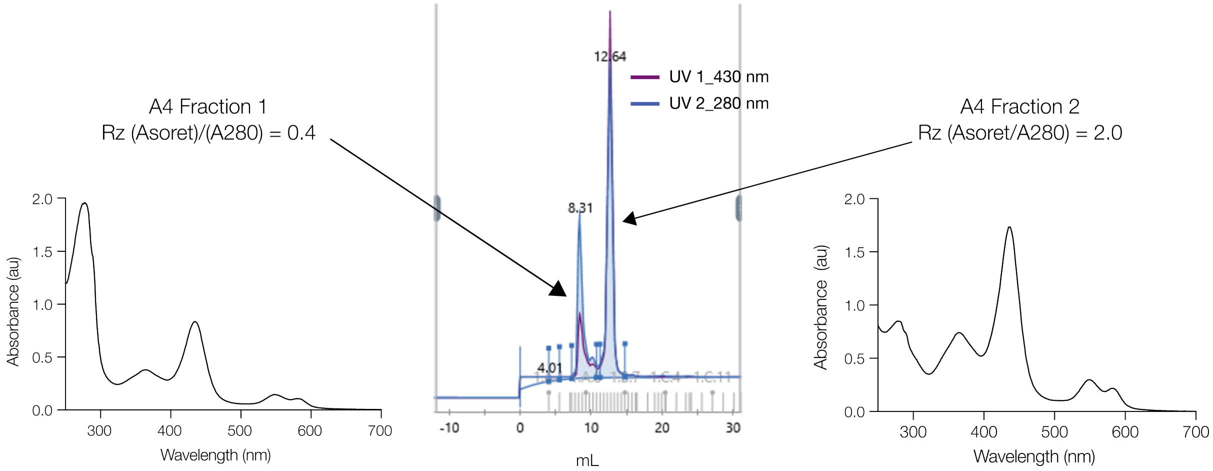

**Supplementary Figure 9.** Separation of fully loaded CoPPIX-A4. Centre: ÄKTA chromatogram from SEC using a Superdex 75 16/600 column (0.6 mL min$^{-1}$) with absorbance monitored at 430 nm (purple) and 280 nm (blue). Two fractions are indicated: left, fraction 1 corresponding to the low-Rz, partially occupied CoPPIX enzyme; right, fraction 2 corresponding to the fully occupied, high-Rz enzyme. Spectra were recorded in a buffer containing 50 mM potassium phosphate, 200 mM NaCl at pH 7.2.

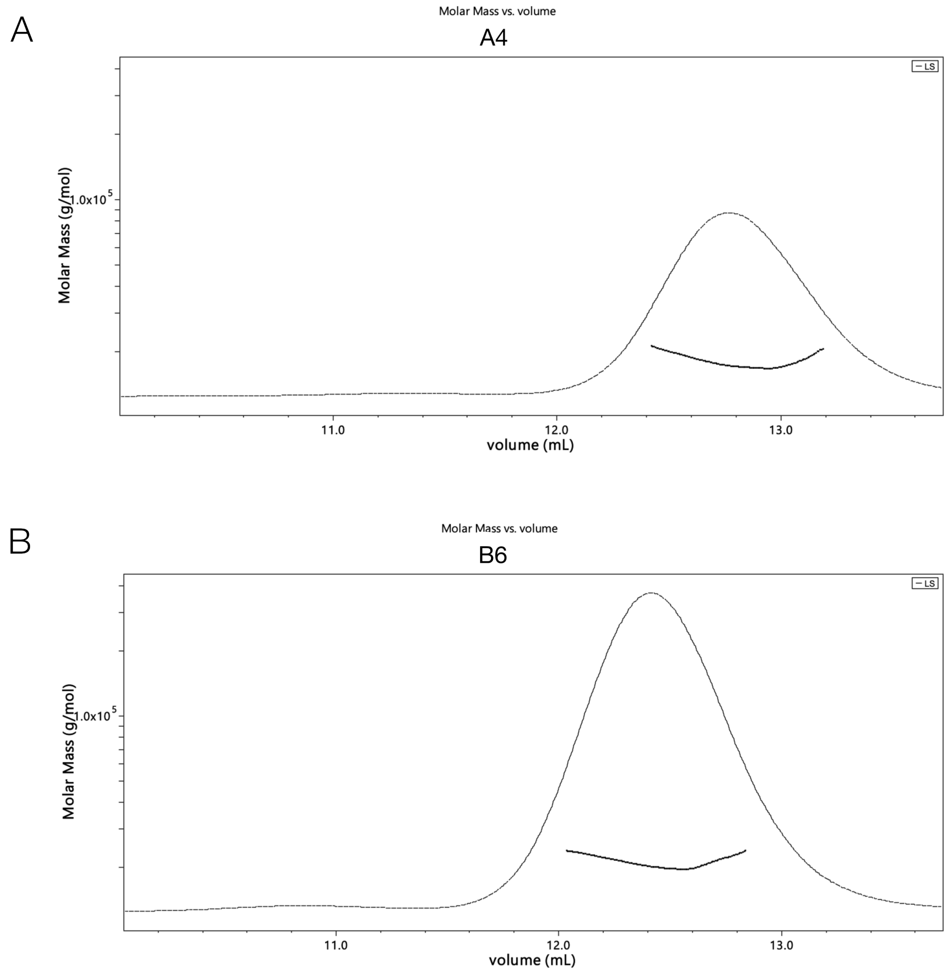

**Supplementary Figure 10.** Elution volume and molar mass determination by SEC-MALS of A4 and B6. Lines represent light scattering and dotted lines represent weight-average molecular mass (MW). Samples elute as monomeric species. A) trace obtained from A4 (1 mg $mL^{-1}$) and elutes as a monomeric species with a MW 18 ± 5.9% kDa. B) trace obtained from B6 (1 mg $mL^{-1)}$ and elutes as a monomeric species with a MW 21 ± 5.9% kDa. Samples run on a Superdex 75 column.

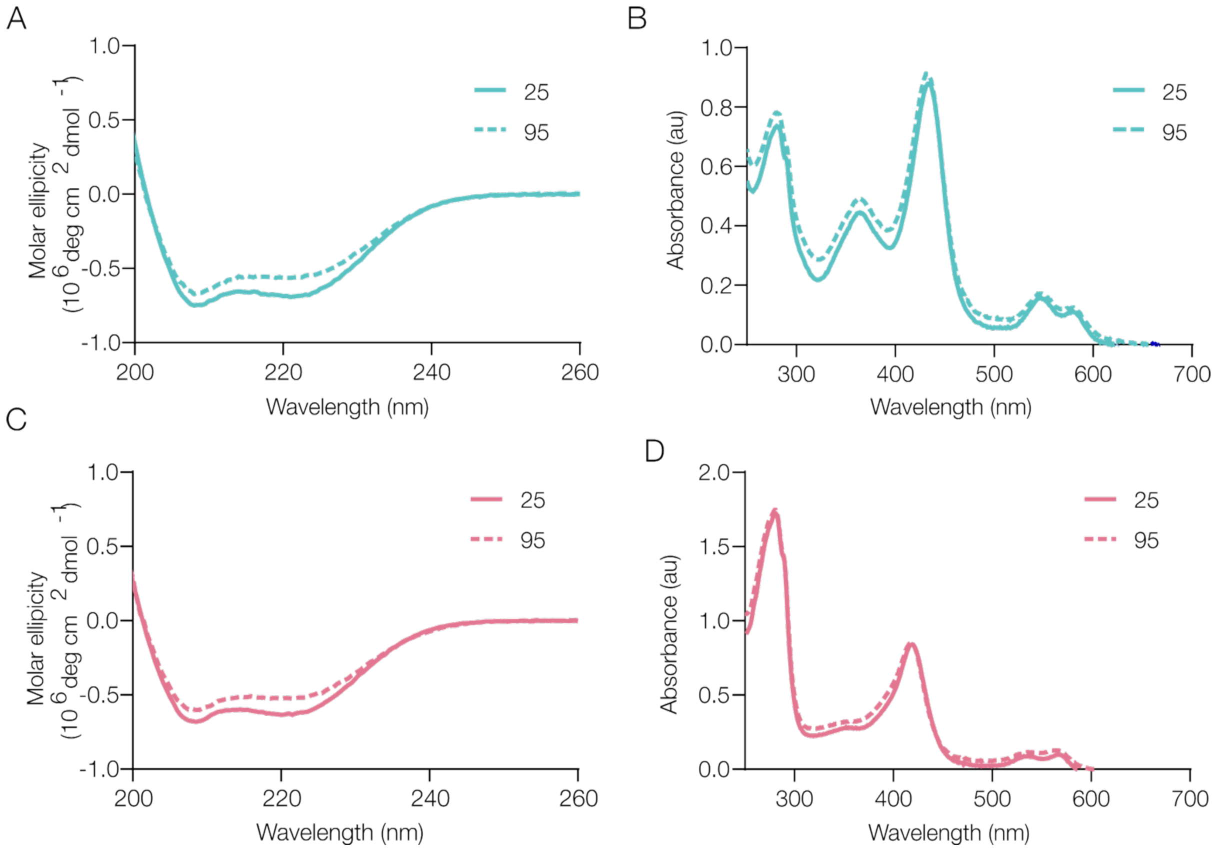

**Supplementary Figure 11.** Thermostability of A4 and B6. A) CD spectra of A4 at 25 °C (solid blue) and 95 °C (dashed blue). B) UV-Vis spectra of A4 at 25 °C (solid blue) and 95 °C (dashed blue) indicating that heme-binding ability is retained at temperatures up to 95 °C. C) CD spectra of B6 at 25 °C (solid pink) and 95 °C (dashed pink). B) UV-Vis spectra of CoPPIX loaded B6 at 25 °C (solid pink) and 95 °C (dashed pink) indicating that heme-binding ability is retained at temperatures up to 95 °C.

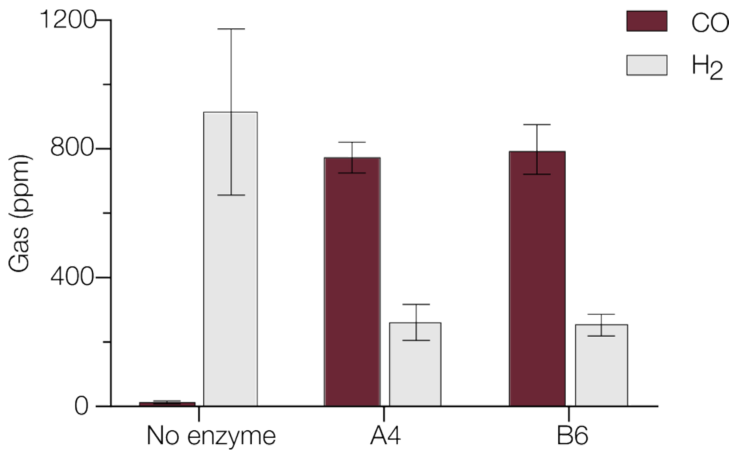

**Supplementary Figure 12.** Activity of no enzyme control, A4 and B6 towards CO (plum) and $H_2$ (grey) in ppm using bicarbonate as the $CO_2$ source. Biotransformations were performed using enzyme (0.3 µM holo-enzyme), 100 mM sodium L-ascorbate, 100mM sodium bicarbonate, and 100 *µ*M $[Ru(bpy)_3]^{2+}$ in 500 mM potassium phosphate buffer (pH 7.0), irradiated under blue light (475 nm) for 30 minutes. A no enzyme control was performed in the absence of enzyme and CoPPIX. Error bars represent the standard deviation of measurements conducted in duplicate.

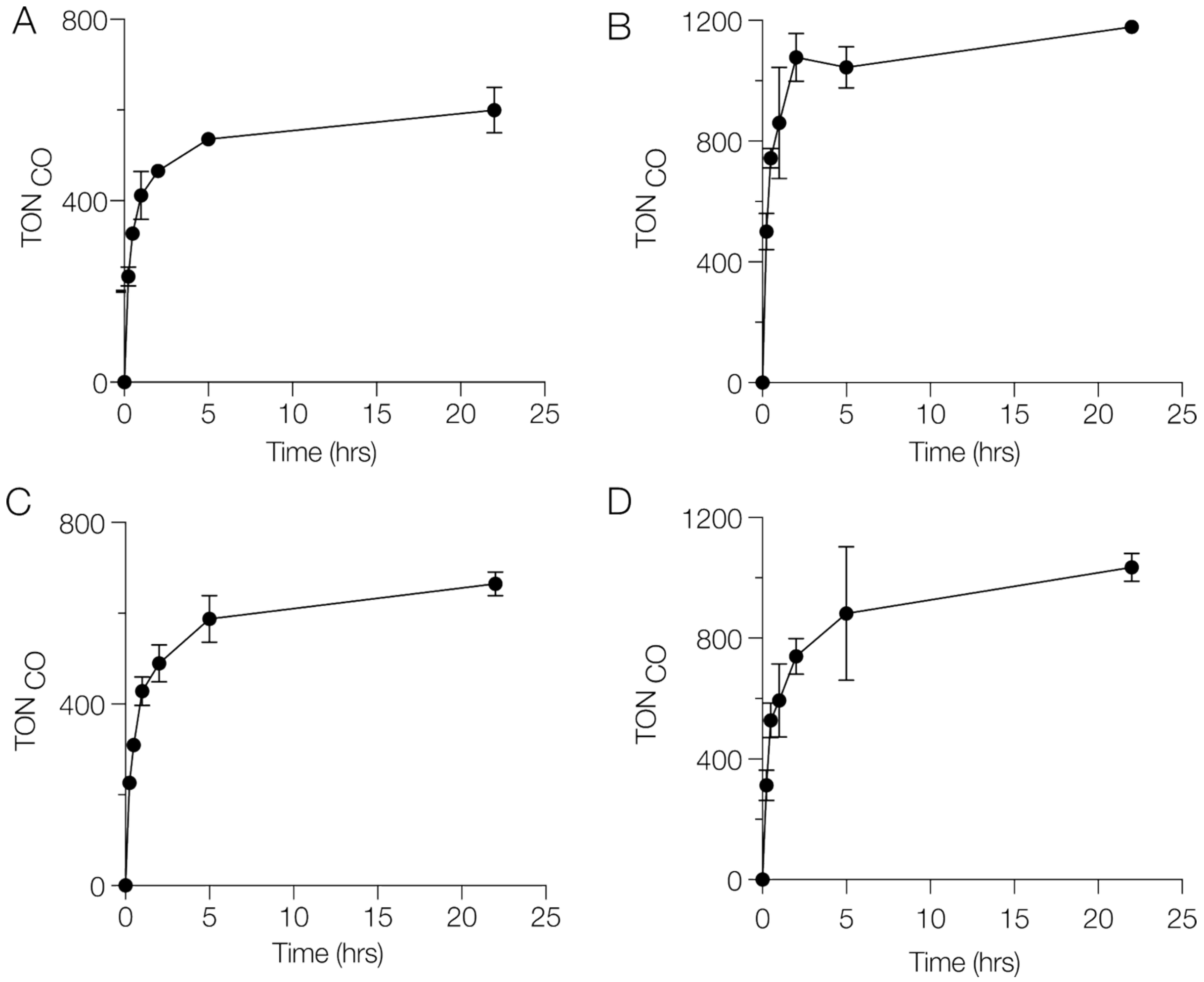

**Supplementary Figure 13.** Time course for CO production during 22 hours of irradiation. A) A4, using bicarbonate as $CO_2$ source. B) A4, using gaseous $CO_2$ C) B6, using bicarbonate as $CO_2$ source. D) B6, using gaseous $CO_2$. Biotransformations were performed using 0.3 µM holoenzyme, 100 mM sodium L-ascorbate, 100mM sodium bicarbonate, 100 $\mu$M $[Ru(bpy)_3]^{2+}$ in 500 mM potassium phosphate buffer (pH 7.0), and irradiated under blue light (475 nm). Error bars represent the range of values from measurements made in duplicate.

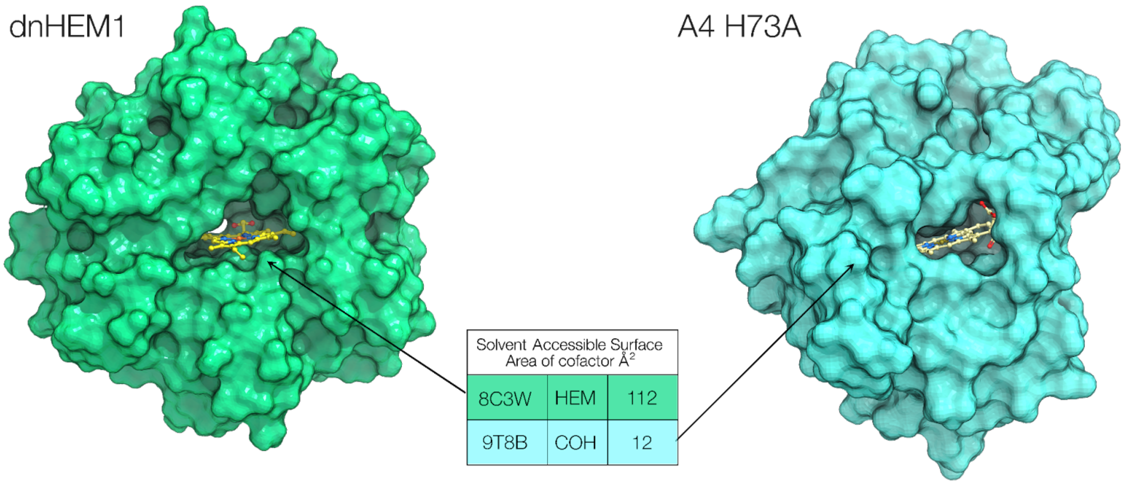

| Solvent Accessible Surface Area of cofactor Å$^2$ | | |
|---|---|---|
| 8C3W | HEM | 112 |
| 9T8B | COH | 12 |

**Supplementary Figure 14.** Surface of dnHEM1 (PDB: 8C3W, green) and A4 H73A (PDB: 9T8B, teal) highlighting the solvent accessible channel to the active sites. CoPPIX cofactor (COH) and heme (HEM) cofactor shown as atom-coloured sticks.

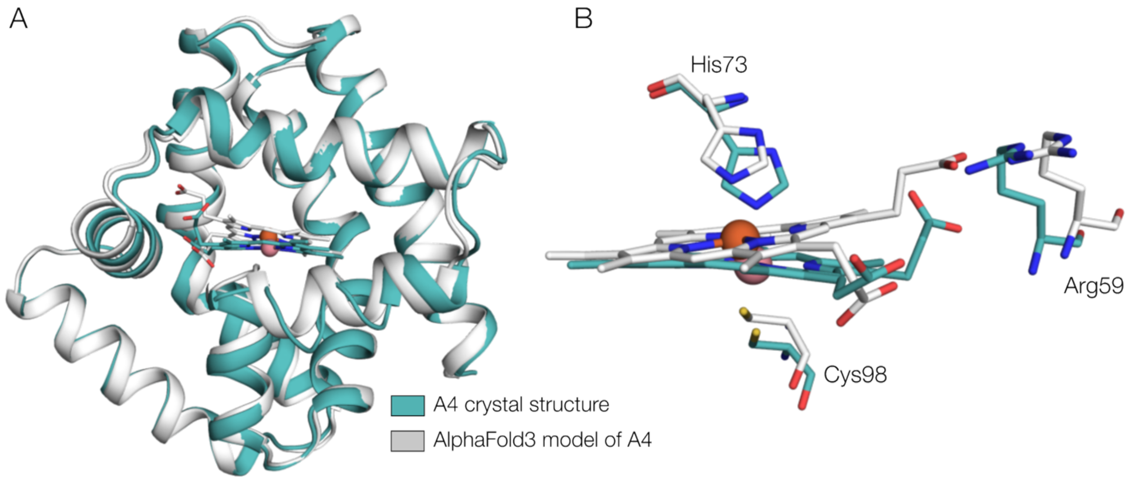

**Supplementary Figure 15.** A) The crystal structure of A4 (PBD: 9T8A, teal) closely matches the AlphaFold3 model (grey). B) Superimposition of the active sites shows the His73 adopts a hexacoordinate position above the CoPPIX cofactor, with additional stabilisation of the propionate group provided by the Arg59, as observed in the predicted AlphaFold3 model.

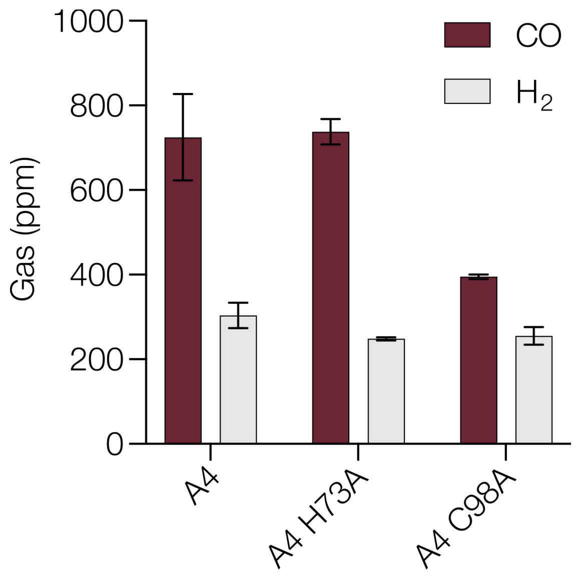

**Supplementary Figure 16.** Photocatalytic $CO_2$ reductase activity of A4 and variants. Bar chart representing the amount of CO (plum) and $H_2$ (grey) in ppm. Biotransformations were performed using 0.3 µM holo-enzyme, 100 mM sodium L-ascorbate, 100 mM sodium bicarbonate, 100 µM $Ru(bpy)^3]^{2+}$ in potassium phosphate buffer (pH 7.0), irradiated with blue light at 475 nm for 30 min. Error bars represent the range of values from measurements made in duplicate.

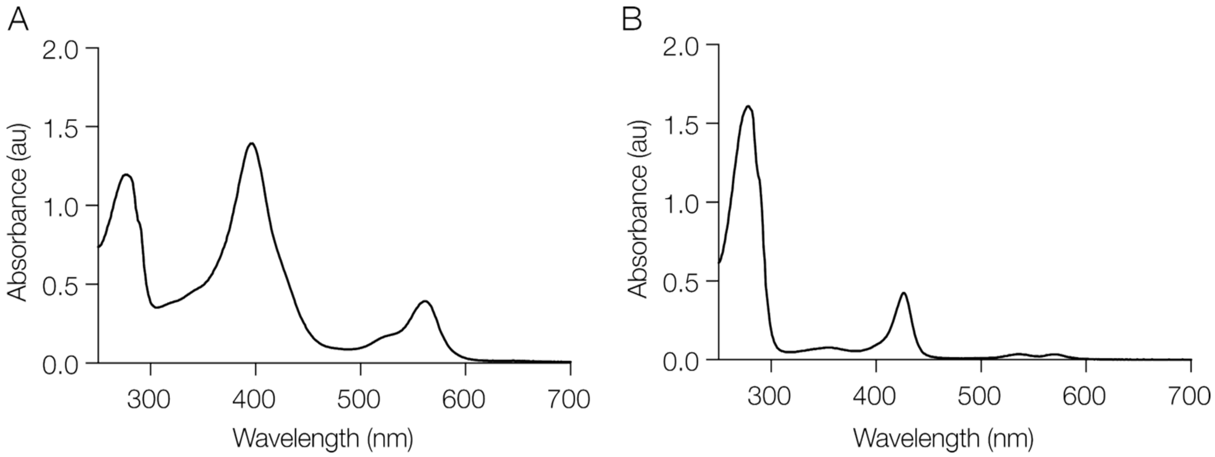

**Supplementary Figure 17.** UV-Vis spectra of A4 variants. A) A4 H73A. B) A4 C98A. Spectra were recorded in a buffer containing 50 mM potassium phosphate, 200 mM NaCl at pH 7.2.

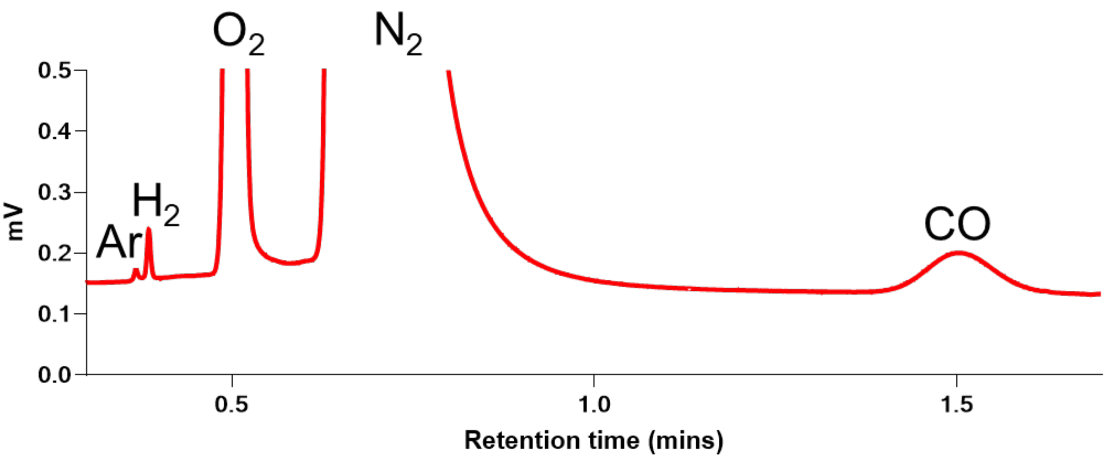

**Supplementary Figure 18.** Micro-GC trace of a reaction using a thermal conductivity detector (TCD). Gases and their respective peaks labelled. Biotransformations were performed using 0.3 µM holo-enzyme, 100 mM sodium L-ascorbate, 100 mM sodium bicarbonate, and 100 µM Ru(bpy)$^{3}$]$^{2+}$ in potassium phosphate buffer (pH 7.0), and irradiated with blue light at 475 nm for 22 hours.

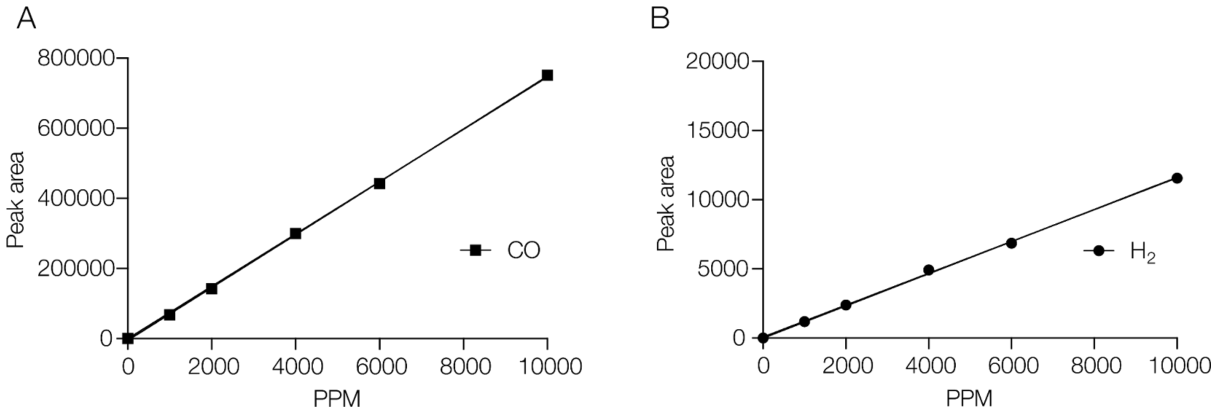

**Supplementary Figure 19.** Gas calibration curves. A) CO *y = 74.615x* B) $H_2$ *y = 1.1623x*. Curves were constructed by injecting known gas standards into a microGC-TCD and plotting the ppm versus observed peak area.

### 2. Supplementary Tables

**Supplementary Table 1.** Design name assignment between heme binder studies and CoPPIX studies.

| Short name | Design name | Axial residue | Design method | Well | Design name | Axial residue | Design method |
|---|---|---|---|---|---|---|---|
| **A1** | HEM_3.A1 * | CYS | RFdiffusion All-Atom | D11 | dnMb11 ** | HIS | ProteinMPNN redesigns (myoglobin) |
| **A2** | HEM_3.A6 | CYS | RFdiffusion All-Atom | D12 | dnMb12 ** | HIS | ProteinMPNN redesigns (myoglobin) |
| **A3** | HEM_3.A8 | CYS | RFdiffusion All-Atom | E1 | dnMb13 ** | HIS | ProteinMPNN redesigns (myoglobin) |
| **A4** | HEM_3.B1 | CYS | RFdiffusion All-Atom | E2 | dnMb14 ** | HIS | ProteinMPNN redesigns (myoglobin) |
| **A5** | HEM_3.B10 | CYS | RFdiffusion All-Atom | E3 | dnMb15 ** | HIS | ProteinMPNN redesigns (myoglobin) |
| **A6** | HEM_3.C7 * | CYS | RFdiffusion All-Atom | E4 | dnMb16 ** | HIS | ProteinMPNN redesigns (myoglobin) |
| **A7** | HEM_3.C9 * | CYS | RFdiffusion All-Atom | E5 | dnMb18 ** | HIS | ProteinMPNN redesigns (myoglobin) |
| **A8** | HEM_3.D4 * | CYS | RFdiffusion All-Atom | E6 | dnMb19 ** | HIS | ProteinMPNN redesigns (myoglobin) |
| **A9** | HEM_3.D9 * | CYS | RFdiffusion All-Atom | E7 | dnMb20 ** | HIS | ProteinMPNN redesigns (myoglobin) |
| **A10** | HEM_3.E5 * | CYS | RFdiffusion All-Atom | E8 | dnMb21 ** | HIS | ProteinMPNN redesigns (myoglobin) |
| **A11** | HEM_3.E6 * | CYS | RFdiffusion All-Atom | F1 | ntf2HEM_2 | HIS | NTF2s |
| **A12** | HEM_3.E7 * | CYS | RFdiffusion All-Atom | F2 | ntf2HEM_17 | HIS | NTF2s |
| **B1** | HEM_3.G1 * | CYS | RFdiffusion All-Atom | F3 | ntf2HEM_19 | HIS | NTF2s |
| **B2** | HEM_3.G2 * | CYS | RFdiffusion All-Atom | F5 | ntf2HEM_25 | CYS | NTF2s |
| **B3** | HEM_3.G7 * | CYS | RFdiffusion All-Atom | F6 | ntf2HEM_15 | CYS | NTF2s |
| **B4** | HEM_2.B4 * | CYS | RFdiffusion All-Atom | F7 | dnHEM1_CYS8.i1_16 | CYS | Inpainting (Cys dnHEM1) |
| **B5** | HEM_2.B11 * | CYS | RFdiffusion All-Atom | F8 | dnHEM1_CYS8.i1_22 | CYS | Inpainting (Cys dnHEM1) |
| **B6** | HEM_1.E6 | CYS | RFdiffusion All-Atom | F9 | dnHEM1_CYS8.i2_11 | CYS | Inpainting (Cys dnHEM1) |
| **B7** | HEM_1.D5 | CYS | RFdiffusion All-Atom | F10 | dnHEM1_CYS8.i2_27 | CYS | Inpainting (Cys dnHEM1) |
| **B8** | HEM_1.D7 * | CYS | RFdiffusion All-Atom | MP.1 | dnHem1.2B.1 | HIS | ProteinMPNN (dnHEM1.2B) |
| **B9** | HEM_2.A6 * | CYS | RFdiffusion All-Atom | MP.3 | dnHem1.2B.3 | HIS | ProteinMPNN (dnHEM1.2B) |
| **B10** | HEM_2.B10 * | CYS | RFdiffusion All-Atom | MP.4 | dnHem1.2B.4 | HIS | ProteinMPNN (dnHEM1.2B) |
| **B11** | HEM_2.C3 * | CYS | RFdiffusion All-Atom | MP.5 | dnHem1.2B.5 | HIS | ProteinMPNN (dnHEM1.2B) |
| **B12** | HEM_2.D10 * | CYS | RFdiffusion All-Atom | MP.6 | dnHem1.2B.6 | HIS | ProteinMPNN (dnHEM1.2B) |
| **D1** | Myoglobin 3RGK | HIS | | MP.7 | dnHem1.2B.7 | HIS | ProteinMPNN (dnHEM1.2B) |

| | | | | | | | |
|---|---|---|---|---|---|---|---|
| **D2** | dnMb2 ** | HIS | ProteinMPNN redesigns (myoglobin) | MP.8 | dnHem1.2B.8 | HIS | ProteinMPNN (dnHEM1.2B) |
| **D3** | dnMb3 ** | HIS | ProteinMPNN redesigns (myoglobin) | MP.9 | dnHem1.2B.9 | HIS | ProteinMPNN (dnHEM1.2B) |
| **D4** | dnMb4 ** | HIS | ProteinMPNN redesigns (myoglobin) | MP.10 | dnHem1.2B.10 | HIS | ProteinMPNN (dnHEM1.2B) |
| **D5** | dnMb5 ** | HIS | ProteinMPNN redesigns (myoglobin) | MP.11 | dnHem1.2B.11 | HIS | ProteinMPNN (dnHEM1.2B) |
| **D6** | dnMb6 ** | HIS | ProteinMPNN redesigns (myoglobin) | Mp.12 | dnHem1.2B.12 | HIS | ProteinMPNN (dnHEM1.2B) |
| **D7** | dnMb7 ** | HIS | ProteinMPNN redesigns (myoglobin) | MP.13 | dnHem1.2B.13 | HIS | ProteinMPNN (dnHEM1.2B) |
| **D8** | dnMb8 ** | HIS | ProteinMPNN redesigns (myoglobin) | MP.14 | dnHem1.2B.14 | HIS | ProteinMPNN (dnHEM1.2B) |
| **D9** | dnMb9 ** | HIS | ProteinMPNN redesigns (myoglobin) | MP.15 | dnHem1.2B.15 | HIS | ProteinMPNN (dnHEM1.2B) |
| **D10** | dnMb10 ** | HIS | ProteinMPNN redesigns (myoglobin) | MP.16 | dnHem1.2B.16 | HIS | ProteinMPNN (dnHEM1.2B) |

** Previously published in: Krishna et al., 2024*[2]
*** Previously published in: Sumida et al., 2024*[3]

**Supplementary Table 2.** Extinction coefficients (ε) of variants calculated using ICP-OES.

| **Sample** | **Extinction coefficient ($mM^{-1}cm^{-1}$)** |
|---|---|
| Co-dnHEM1 | $\varepsilon_{422}$ 86.9 |
| A4 | $\varepsilon_{434}$ 101.4 |
| B6 | $\varepsilon_{422}$ 97.4 |

**Supplementary Table 3.** Selectivity and turnover numbers of A4 and B6 $CO_2$ biotransformations.

| Enzyme | Concentration | Time | $CO_2$ source | Turnovers | Selectivity % |
|---|---|---|---|---|---|
| A4 | 1 µM | 30 min | Bicarbonate | 195 | 72 |
| A4 | 1 µM | 22 hr | Bicarbonate | 380 | 54 |
| A4 | 0.3 µM | 30 min | Bicarbonate | 396 | 75 |
| A4 | 0.3 µM | 22 hr | Bicarbonate | 599 | 34 |
| A4 | 0.3 µM | 30 min | Gaseous $CO_2$ | 743 | 84 |
| A4 | 0.3 µM | 22 hr | Gaseous $CO_2$ | 1179 | 50 |
| B6 | 1 µM | 30 min | Bicarbonate | 124 | 71 |
| B6 | 1 µM | 22 hr | Bicarbonate | 351 | 48 |
| B6 | 0.3 µM | 30 min | Bicarbonate | 382 | 76 |
| B6 | 0.3 µM | 22 hr | Bicarbonate | 665 | 30 |
| B6 | 0.3 µM | 30 min | Gaseous $CO_2$ | 528 | 84 |
| B6 | 0.3 µM | 22 hr | Gaseous $CO_2$ | 1035 | 40 |

**Supplementary Table 4.** Amino acid and DNA sequences (encoding region and purification tags) of characterized *de novo* designed heme-binding proteins.

| **Design** | **Amino acid sequence**<br><br>**DNA sequence** |
|---|---|
| dnHEM1 | MVSLDQAILILVVAAKLGTTVEEAVKRALWLKTKLGVSLDQALRILSAAANTGTTVEEAVKRALKLKTKLGVSLEAALAILSAAAQLGTTVEEAVK<br>RALKLKTKLGVDLETAALALLTAAKLGTTVEEAVKRALKLKTKLGVSLIEALHILLTAAVLGTTVEEAVYRALKLKTKLGVSLLQAAAILILAARLGTT<br>VEEAVKRALKLKTKLGGGSGGSHHWGSGSHHHHHH<br><br>ATGGTGAGCCTGGATCAGGCGATTCTGATTCTGGTGGTGGCGGCGAAACTGGGCACCACCGTGGAAGAAGCGGTGAAACGCGCGCTG<br>TGGCTGAAAACCAAATTAGGCGTGTCGTTGGACCAGGCGCTGCGTATTCTGAGCGCGGCCGCCAATACCGGCACGACGGTTGAAGAGG<br>CCGTTAAACGTGCACTGAAACTGAAGACGAAGTTGGGTGTTTCGCTGGAAGCGGCGCTGGCGATTTTAAGCGCCGCGGCGCAGCTGG<br>GTACGACCGTTGAGGAGGCGGTTAAGCGCGCGTTGAAATTGAAAACGAAATTGGGGGTGGATCTGGAAACCGCGGCCCTGGCGTTGTT<br>GACCGCAGCCAAGCTCGGTACCACTGTGGAGGAAGCAGTCAAGCGTGCCCTGAAGTTAAAGACCAAGCTGGGGGTCAGCTTGATTGAG<br>GCACTGCATATTCTGCTGACCGCTGCGGTGTTGGGCACTACCGTAGAGGAAGCAGTGTATCGCGCCTTGAAGCTCAAGACTAAGTTAGG<br>TGTTAGTCTGCTGCAGGCGGCAGCCATCTTGATTTTAGCCGCGCGCCTGGGGACGACTGTCGAAGAGGCTGTGAAGCGCGCGCTCAA<br>GTTGAAGACCAAACTCGGTGGCGGGAGCGGTGGCTCTCATCATTGGGGCAGTGGCTCGCATCATCACCACCATCAT |
| dnHEM1.2B | MVSLDQAILILRVAAKLGTTVEEAVKRALWLKTKLGVSLHQALRILSRAANTGTTVEEAVKRALKLKTKLGVSLEAALAILSAAAQLGTTVEEAVK<br>RALKLKTKLGVDLETAALALLTAAKLGTTVEEAVKRALKLKTKLGVSLIEALHILLTAAVLGTTVEEAVYRALKLKTKLGVSLLQAAAILILAARLGTT<br>VEEAVKRALKLKTKLGGGSGGSHHWGSGSHHHHHH<br><br>ATGGTGAGCCTGGATCAGGCGATTCTGATTCTGCGGGTGGCGGCGAAACTGGGCACCACCGTGGAAGAAGCGGTGAAACGCGCGCTG<br>TGGCTGAAAACCAAATTAGGCGTGTCGTTGCACCAGGCGCTGCGTATTCTGAGCCGGGCCGCCAATACCGGCACGACGGTTGAAGAGG<br>CCGTTAAACGTGCACTGAAACTGAAGACGAAGCTCGGTGTTTCGTTAGAGGCGGCGCTGGCGATTTTAAGCGCAGCCGCGCAGCTGGG<br>TACTACTGTGGAGGAGGCGGTTAAGCGCGCGTTGAAATTGAAAACGAAGTTGGGCGTGGATCTGGAAACCGCGGCCTTAGCGTTGTTG<br>ACCGCAGCCAAGTTAGGTACGACCGTTGAGGAAGCAGTTAAGCGCGCCCTGAAGTTAAAGACCAAGTTGGGTGTGAGCTTGATTGAGG<br>CACTGCATATTCTGCTGACTGCCGCGGTGTTAGGCACTACCGTCGAAGAGGCGGTGTATCGCGCCTTGAAGTTGAAAACTAAATTGGGG<br>GTTAGTCTGCTGCAGGCGGCTGCCATCTTGATTTTAGCAGCCCGCCTGGGGACTACGGTGGAGGAGGCCGTAAAGCGTGCCTTAAAATT<br>AAAAACCAAATTGGGTGGGGGCAGCGGTGGCAGCCATCATTGGGGCTCGGGCTCGCATCATCACCACCATCAT |

**Supplementary Table 5.** Mass spectrometry data for variants. (+Met) denotes that the N-terminal methionine is present in the observed mass.

| Construct | Estimated mass (no methionine) | Observed mass |
|---|---|---|
| dnHEM1 | 24030 | 24028 |
| dnHEM1.2B | 24194 | 24193 |
| A4 | 19218 | 19219 |
| A4 (no snac tag) | 18368 | 18368 |
| A4 H73A | 19152 | 19152 |
| B6 | 22591 | 22592 |
| B6 (no snac tag) | 21741 | 21742 |
| HEM_1.D5 **(B7)** | 19885.6 | 19885 |
| HEM_1.E6 **(B6)** | 22592.1 | 22591 |
| HEM_3.A6 **(A2)** | 24642.9 | 24642 |
| HEM_3.A8 **(A3)** | 24998.3 | 24997 |
| HEM_3.B1 **(A4)** | 19219 | 19218 |
| HEM_3.B10 **(A5)** | 23168.4 | 23168 |
| ntf2HEM_2 **(F1)** | 16296.2 | 16295 |
| ntf2HEM_15 **(F6)** | 15230 | 15229 |
| ntf2HEM_17 **(F2)** | 15361.9 | 15492 |
| ntf2HEM_19 **(F3)** | 14804.3 | 14803, 14934 (+Met) |
| ntf2HEM_25 **(F5)** | 15435.3 | 15434 |
| ntf2HEM_2_H53A | 16230.1 | 16229 |
| ntf2HEM_17_H51A | 15295.9 | 15295, 15426 (+Met) |
| ntf2HEM_19_H58A | 14738.2 | 14737, 14868 (+Met) |
| ntf2HEM_19_H35A | 14738.2 | 14737, 14869 (+Met) |
| ntf2HEM_25_C55A | 15403.3 | 15402 |
| dnHEM1_CYS8 | 27095 | 27094 |
| dnHEM1_CYS8.i1_16 **(F7)** | 31721.8 | 31720 |
| dnHEM1_CYS8.i1_22 **(F8)** | 31768.3 | 31767 |
| dnHEM1_CYS8.i2_11 **(F9)** | 31317.9 | 31316 |
| dnHEM1_CYS8.i2_27 **(F10)** | 32813.2 | 32812 |
| dnHEM1.2B.1 **(MP.1)** | 24412 | 24411 |
| dnHEM1.2B.3 **(MP.3)** | 24131 | 24129 |
| dnHEM1.2B.4 **(MP.4)** | 24028 | 24026 |
| dnHEM1.2B.5 **(MP.5)** | 24113 | 24111 |
| dnHEM1.2B.6 **(MP.6)** | 24103 | 24101 |

| | | |
|---|---|---|
| dnHEM1.2B.7 **(MP.7)** | 24165 | 24163 |
| dnHEM1.2B.8 **(MP.8)** | 24550 | 24549 |
| dnHEM1.2B.9 **(MP.9)** | 24160 | 24158 |
| dnHEM1.2B.10 **(MP.10)** | 24561 | 24560 |
| dnHEM1.2B.11 **(MP.11)** | 24001 | 23999 |
| dnHEM1.2B.12 **(MP.12)** | 24166 | 24164 |
| dnHEM1.2B.13 **(MP.13)** | 23967 | 23965 |
| dnHEM1.2B.14 **(MP.14)** | 24290 | 24288 |
| dnHEM1.2B.15 **(MP.15)** | 24006 | 24005 |
| dnHEM1.2B.16 **(MP.16)** | 24571 | 24569 |

**Supplementary Table 6.** Data collection and refinement statistics of A4 and A4 H73A. Data for the outermost resolution shell are given in parenthesis. R-free was calculated using ~5% of the data separate from the rest.

| | **A4** | **A4 H73A** |
|---|---|---|
| Wavelength (Å) | 0.9769 | 0.9763 |
| PDB ascension number | 9T8A | 9T8B |
| Resolution range | 71.8 - 2.08<br>(2.14 - 2.08) | 45.09 - 1.81<br>(1.88 - 1.81) |
| Space group | P 1 21 1 | P 1 21 1 |
| Unit cell<br>a, b, c, (Å)<br>α, β, γ (°) | 44.212 161.742 74.587<br>90 105.72 90 | 45.101 63.427 51.477<br>90 88.61 90 |
| Total reflections | 477737 (36325) | 153934 (17161) |
| Unique reflections | 67786 (5158) | 22256 (2454) |
| Multiplicity | 7.0 (7.0) | 6.9 (7.0) |
| Completeness (%) | 99.92 (99.85) | 99.89 (99.47) |
| Mean I/sigma(I) | 7.40 (0.62) | 18.74 (2.25) |
| Wilson B-factor | 34.83 | 36.07 |
| R-merge | 0.1412 (1.531) | 0.05026 (0.582) |
| R-meas | 0.1524 (1.652) | 0.0543 (0.6289) |
| R-pim | 0.05699 (0.6182) | 0.02038 (0.2366) |
| CC1/2 | 0.998 (0.559) | 0.999 (0.882) |
| CC* | 0.999 (0.847) | 1 (0.968) |
| Reflections used in refinement | 60270 (4609) | 22241 (2451) |
| Reflections used for R-free | 1782 (120) | 1114 (139) |
| R-work | 0.1990 (0.3063) | 0.1940 (0.2571) |
| R-free | 0.2355 (0.3532) | 0.2046 (0.2494) |
| Number of non-hydrogen atoms | 7983 | 2650 |
| macromolecules | 7289 | 2429 |
| ligands | 348 | 143 |
| solvent | 346 | 78 |
| Protein residues | 936 | 315 |
| RMS (bonds) | 0.012 | 0.009 |
| RMS (angles) | 1.12 | 0.87 |
| Ramachandran favored (%) | 98.05 | 98.07 |

| | | |
|---|---|---|
| Ramachandran allowed (%) | 1.84 | 1.93 |
| Ramachandran outliers (%) | 0.11 | 0 |
| Rotamer outliers (%) | 0.38 | 0.38 |
| Clashscore | 8.26 | 5.31 |
| Average B-factor | 44.14 | 45.32 |
| macromolecules | 44.38 | 45.01 |
| ligands | 39.68 | 49.79 |
| solvent | 43.57 | 46.9 |

### 3. De novo Heme-binder library design and characterization

#### 3.1. Methods and Materials

***General protocol for cloning and testing designed proteins for heme binding.***

The amino acid sequences of designed proteins were reverse translated to corresponding DNA sequences, optimized for synthesizability and expressibility. The synthetic DNA fragments encoding the designed proteins were obtained from IDT as eBlocks. The DNA fragments contained overhangs (5’ atactacggtctcaagga- and -ggttcccgagaccgtaatgc 3’) for Golden Gate cloning. The DNA fragments were cloned into LM627 vector (Addgene #191551), encoding C-terminal SNAC and 6His tags for purification, with the final expressed sequence being MSG-<design>-GSGSHHWGSTHHHHHH.

The cloned DNA vectors were transformed into *E. coli* BL21(DE3) competent cells (NEB C2527I) and grown up in SOC media at 37 °C. 1 mL LB-kanamycin cultures were grown in a 96-well round-bottom deep-well plate (37 °C for 18 hours, shaking at 1,100 r.p.m.). 100 µL of the LB culture was added to 900 µL of Terrific Broth II + kanamycin in 96-well round-bottom deep-well plate, the cultures were grown at 37 °C for 3 hours in a shaking incubator at 1,100 r.p.m., and thereafter 1 mM of IPTG was added to induce protein expression. The plates were further incubated for 4 hours, after which the cells were collected using centrifugation and frozen at -20 °C.

#### 3.2. Redesign of dnHEM1.2B with ProteinMPNN

The input structures for redesign were obtained from AlphaFold2[4] predictions of dnHEM1.2B. The structure was predicted with three AF2 models (models 3, 4, 5), then the heme ferryl intermediate was placed in the pocket and the structure was relaxed with Rosetta FastRelax[5] 5 separate times. This nudged the ligand more into place and readjusted some rotamers and the backbone. This approach yielded 15 backbones for the same sequence that slightly differed from each other.

To design new sequences for dnHEM1.2B, ProteinMPNN[6] was used at four different temperatures : 0.1, 0.2, 0.5, 0.7, with 5 sequences generated at each temperature for each backbone. 300 sequences were created in total across all of the 15 input structures. During sequence generation, all of the positions in the immediate vicinity of heme were kept fixed. Those positions were: 4, 7, 8, 11, 15, 39, 40, 43, 46, 47, 50, 74, 78, 81, 85, 107, 109, 110, 112, 113, 116, 120, 144, 148, 151, 155, 177, 179, 180, 183, 186, 190.

Thereafter, the quality of the sequences was evaluated with AlphaFold2 (model 4, 3 recycles). Sequences were selected based on the following metrics: plDDT > 90.0, $C_\alpha$ RMSD < 0.7, HIS sidechain RMSD < 0.7. As a result, 8 passing sequences were obtained from ProteinMPNN T=0.5/0.7, and 40 passing designs from T=0.1/0.2. All eight sequences from the higher temperature set were selected for characterization. From the lower temperature set, eight sequences were picked based on sequence similarity clustering.

#### 3.3. Design of heme-binding proteins in NTF2 fold

**Computational design**

To design a heme binding site into the nuclear transport factor 2 (NTF2) protein scaffold, we used an analogous design strategy (Rosetta Matcher and FastDesign; described below) as was used to design the dnHEM1 heme-binding protein[7]. We used the heme model with a Fe-bound methyl group in the distal site, and a ligating histidine or cysteine residue in the proximal side. The NTF2 protein scaffolds were generated with a TrRosetta-based hallucination method using constraints derived from NTF2-like structures and homology models including sequence and geometric constraints for recapitulation of native hydrogen-bonding networks[8,9]. The foldability of the parent scaffold sequences were verified with AlphaFold2. Rosetta Matcher was used to geometrically place the coordinating histidine or cysteine residue and the heme ligand into the pockets of these protein scaffolds, identifying suitable positions that can host the histidine/cysteine inverse rotamer while not introducing clashes between the heme and the protein backbone. In each of the scaffolds the pre-installed hydrogen bond network positions were annotated and omitted from the matching procedure. The amino acid sequence at positions surrounding the heme and the histidine were optimized using Rosetta FastDesign[10]. During the design, constraints were applied to the heme-HIS/CYS interaction, and the residue identities in the native H-bond network were kept fixed. After the design step, the structures were relaxed without the applied constraints, and each model was scored for metrics describing the protein-ligand interaction (H-bonds to COO groups, SASA, contact molecular surface), the quality of the native H-bond network, and the coordinating histidine/cysteine rotamer quality and constraint score. Passing designs were lastly evaluated with AlphaFold2 for how well the predicted structure recapitulates the design model. The predictions were performed with single sequence input, using model 4 and 20 recycles. Promising designs were selected based on confidence (plDDT > 90.0), backbone accuracy ($C_\alpha$ RMSD < 1.0 Å) and coordinating histidine/cysteine accuracy (side chain RMSD < 1.2 Å).

**Experimental characterization**

Cloning and small scale *E. coli* culture preparation was performed following the "general protocol for cloning and testing designed proteins for heme binding".

The small cultures were lysed to determine which variants were overexpressed and soluble, and to qualitatively determine their heme-binding abilities. Lysis was performed using a 9:1 mixture of TBS (25 mM Tris, 300 mM NaCl, pH 8) and BugBuster (10X) lysis reagent, containing 10 µg/mL DNAse I, using microplate sonication bath (Sonica Q500 with a microplate horn) with a total of 10 minutes "on" time at 40% amplitude. 35/50 designs were found to give soluble expression, as judged by SDS-PAGE analysis. To determine heme-binding of variants with soluble expression, 3 µM hemin was added to the clarified lysates and UV-Vis spectra were recorded. Promising heme-binders were detected based on a notable Soret band features in the 400-415 nm range. 15 designs were deemed as promising heme-binders based on this analysis and their expression was scaled up for more detailed studies.

Fifteen promising designs were grown up as 50 mL cultures, isolated in the presence of heme (added to lysis buffer before sonication), and purified with size-exclusion chromatography. The heme-binding of the designed proteins in NTF2 scaffold was characterized by UV-Vis spectroscopy. 10 µM of purified protein was mixed with 2 µM hemin and spectra were recorded (Supplementary Figure 20). Good designs were identified based on the presence of a sharp and intense Soret band (at 400-410 nm region). Six designs were found to have good Soret features, and express in good yield and purify as a single oligomer. Variants bearing an alanine mutation at the putative heme-coordinating residue of promising designs were expressed and UV-Vis spectra of 10 µM purified protein with 2 µM hemin was measured. In all cases we observed spectral changes compared to the parent design.

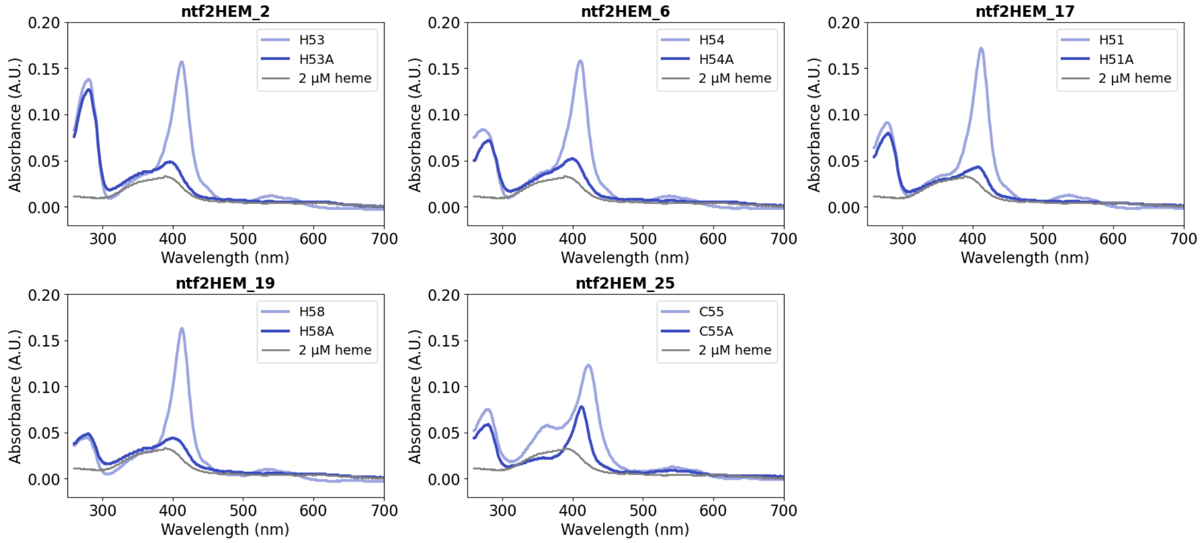

**Supplementary Figure 20.** UV-Vis spectra of purified heme-binding residue mutants (HIS or CYS to ALA) of designed heme binding proteins in NTF2 scaffold. 10 µM protein was mixed with 2 µM hemin. The gray spectrum line corresponds to 2 µM hemin in TBS buffer.

#### 3.4. Design of CYS-based heme-binding proteins with inpainting

To convert the histidine-based heme-binding protein dnHEM1 into a cysteine-based heme-binding protein we placed a new CYS-containing backbone motif (from cytochrome P450; PDB: 2QBL[11]) underneath the heme using PyMol. We used RFjoint2 inpainting[12] to connect the CYS-motif with the rest of the protein scaffold, remodelling the adjacent loops and helices. Since RFjoint2 model is unable to see the presence of ligands, a triphenylalanine peptide chain was placed into the heme binding site to mimic the steric presence of heme. Thereafter, we used ProteinMPNN to generate amino acid sequences at positions that were built with RFjoint2 inpainting and used AlphaFold2 (single sequence, model 4, 3 recycles) to identify promising scaffolds for heme binding site redesign. For heme binding site design, heme model containing a methyl group in the distal site was aligned into the AlphaFold2 predicted models from dnHEM1. Rosetta FastDesign and ProteinMPNN were used to optimize the amino acid sequence surrounding the heme ligand, adjusting hydrophobic contacts and creating H-bond interactions. The design models were scored for metrics describing the protein-ligand interaction (H-bonds to COO groups, SASA, contact molecular surface), the coordinating histidine rotamer quality and constraint score. The designed sequences were again predicted with AlphaFold2 and filtered based on using confidence and accuracy metrics (plDDT > 87.0, $C_\alpha$ RMSD < 1.2 Å and CYS RMSD < 1.0 Å). Twenty-five designs were selected for experimental characterization.

The synthetic DNA encoding these designs was obtained from IDT, and cloned into LM627 vector (AddGene #191551) using Golden Gate Assembly. The plasmid was transformed into *E. coli* BL21(D3) cells and 40 mL cultures in TB-II-kanamycin were grown of each variant. Proteins were isolated from cells by ultrasonication in the presence of excess heme, and the *holo* proteins were isolated by Ni-NTA IMAC and size-exclusion chromatography. While most proteins retained some amount of heme and displayed some Soret band in the UV-Vis spectra, one variant in particular (dnHEM1-CYS8) had a strong heme signal with a distinctive 420 nm peak corresponding to CYS-ligated heme. This variant was characterized further, confirming its high thermostability by variable temperature UV-Vis and circular dichroism, and it's heme binding was CYS-dependent as judged by significant changes in the UV-Vis spectra upon C/A mutation (Supplementary Figure 21).

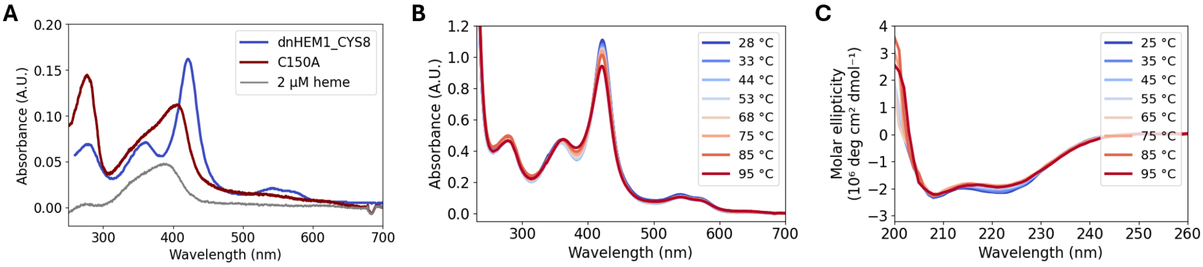

**Supplementary Figure 21.** Spectral characterization of dnHEM1-CYS8. A) UV-Vis spectrum of C150A mutant of dnHEM1-CYS8. B) UV-Vis spectra collected at increasing temperatures. C) Circular dichroism (CD) spectra collected at increasing temperatures.

Having confirmed that dnHEM1-CYS8 is a good heme-binding protein, we used this design as a starting point for further diversification of the protein scaffold and potential substrate binding site with RFjoint2 inpainting. We followed two diversification strategies: (1) opening up the top of the heme binding site and closing the front; (2) opening up the top, closing the front and replacing the P450 CYS-motif with a motif from unspecific peroxygenase (Supplementary Figure 22).

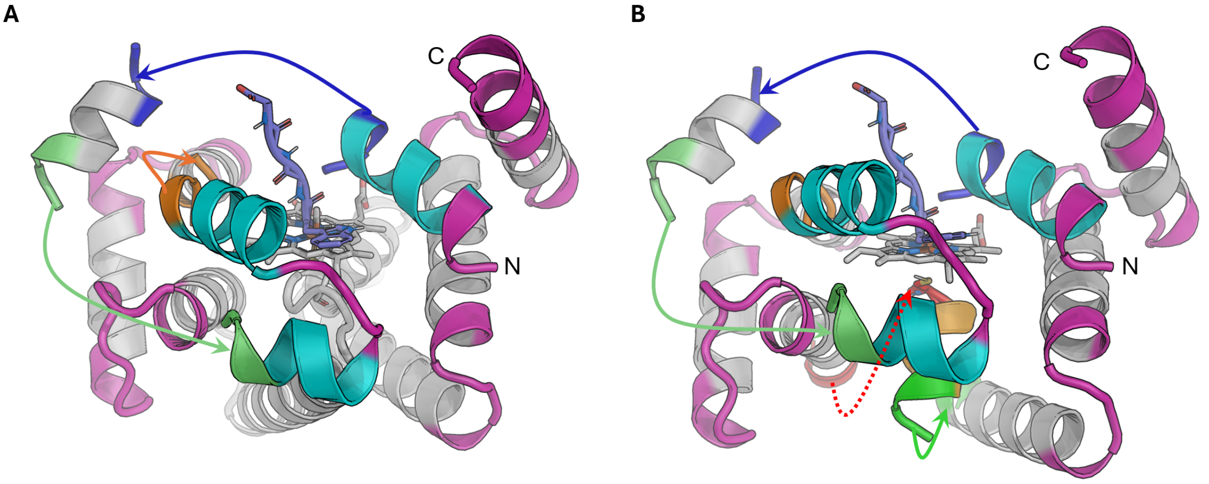

**Supplementary Figure 22.** RFjoint2 input structure for converting dnHEM1-CYS8 from front-open to top-open heme-binding protein. Gray elements were fixed from the parent scaffold, and magenta parts were remodelled in the context of new connections. Cyan elements are newly added guiding elements. New connections are shown with coloured arrows. Purple peptide in the middle is guiding the substrate and heme binding sites. A) Strategy 1: keeping the CYS motif from dnHEM1-CYS. B) Strategy 2: changing the CYS motif to that from an unspecific peroxygenase (orange helix, PDB: 7O2G[13]).

**Diversification strategy 1: opening up the top and closing the front**

The design model (AlphaFold2 predicted backbone, relaxed with heme) of dnHEM1-CYS8 was used as a starting point. To create an opening at the top we took inspiration from the native unspecific peroxygenase protein family where the substrate binding site above heme is lined by two parallel helices. To achieve that in dnHEM1-CYS8, we removed the top helices, extended the left helix out and connected with a helix-loop-helix motif that blocks the heme binding site from the front. On the right side, the guiding helix was placed in the opposite direction to the helix that was there in dnHEM1, antiparallel to the left side helix. A small helix was kept in the top-left corner to guide the backbone generation. A small polypeptide with sequence WGGG was placed in the heme binding site and between the helices to leave space for substrate entry and for the heme. RFjoint2 inpainting was used to generate backbones from these guiding secondary structure elements, rebuilding 10-50 amino acid fragments to join them.

**Diversification strategy 2: opening up the top, closing the front, using UPO-CYS motif**

The design model (AlphaFold2 predicted backbone, relaxed with heme) of dnHEM1-CYS8 was used as a starting point to create an opening at the top and close up the front. The CYS-motif from unspecific peroxygenase (PDB: 7O2G[13]) was placed underneath the heme, replacing the existing P450-like CYS motif. This motif only fits into this scaffold with its C-terminus angled forward (as opposed to the P450 motif which has the C-terminus towards the back). For that reason, the bottom helix was removed, and various connection strategies were explored to connect the motif with the rest of the scaffold. RFjoint2 inpainting was used to generate backbones from these guiding secondary structure elements, rebuilding 10-50 amino acid fragments to join them.

The same design workflow as before in the design of dnHEM1-CYS8 was used to turn the designed backbones into functional designs. ProteinMPNN and AlphaFold2 were used to design the initial sequence onto the entire backbone, and identify well-folding sequences (plDDT > 87.0, $C_\alpha$ RMSD < 1.5 Å, CYS RMSD < 1.2 Å). To design the heme binding site, along with a generic substrate, a heme model with *para*-tolyl group in the distal site was used. This ensured a larger hydrophobic opening, compared to the methyl group used before. Rosetta FastDesign was used to design sequences around the heme binding site, and LigandMPNN[14] was used to generate additional sequence diversity in the second shell. Finally, AlphaFold2 (single sequence, model 4, 3 recycles) was used to identify final well-folding design candidates (plDDT > 90.0, $C_\alpha$ RMSD < 1.0 Å and CYS RMSD < 0.8 Å). Twenty-three designs were selected from strategy 1, and twenty-seven from strategy 2 for experimental testing, and their characterization is described below.

**Experimental characterization**

Cloning and small-scale *E. coli* culture preparation was performed following the "general protocol for cloning and testing designed proteins for heme binding".

The LB culture was added to 40 mL of Terrific Broth II + kanamycin autoinduction media, the cultures were grown at 37 °C for 18 hours. Small aliquots were taken from the cultures for SDS-PAGE analysis and the rest of the cells harvested by centrifugation and frozen at -20 °C. The small aliquots of cells were lysed using BugBuster lysis reagent, together with 10 µg/mL DNAse I, and clarified by centrifugation. The supernatant was analyzed by SDS-PAGE. 37 designs were found to give soluble overexpression - 16/24 from strategy 1 and 21/28 from strategy 2.

Thawed cell pellets of well expressing designs were lysed in the presence of 300 µM hemin, the protein isolated by Ni-NTA IMAC, and purified by size-exclusion chromatography. The purified heme-loaded proteins were analyzed with UV-Vis spectrophotometry (Jasco Spec V750, 1 cm pathlength) to characterize their heme binding properties (Supplementary Figure 23 & 24). The spectra collected at room temperature indicated that all proteins contained heme to some degree, showing prominent Soret features corresponding to heme-CYS ligation with heme in low spin hexacoordinate state ($\lambda_{Soret}$ ~420 nm) or in high spin pentacoordinate state ($\lambda_{Soret}$ ~385 nm). Heating the protein samples up to 95 °C and collecting UV-Vis spectra at every 10 °C showed that most of the design proteins retain their heme-binding ability (Supplementary Figure 25). A few variants undergo spin-state change, potentially due to the unbinding of a distal residue or an imidazole molecule.

Taken together this campaign uncovered a number of well-behaving inpainted CYS-containing proteins that bind heme. We selected four variants into the heme-binder library for $CO_2$ reductase activity testing.

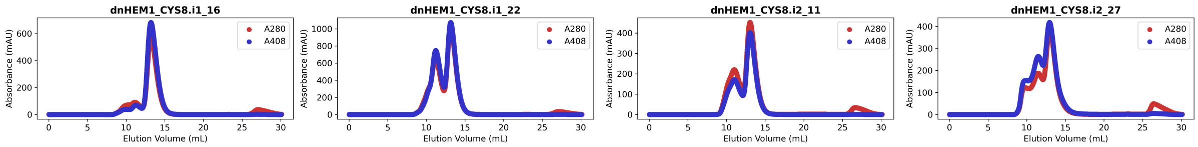

**Supplementary Figure 23.** Size-exclusion chromatograms of purified heme-binding proteins, using Superdex Increase 75 10/300 GL column and 25 mM Tris 300 mM NaCl pH 8 running buffer. Red trace corresponds to a chromatogram based on absorption at 280 nm, blue trace is collected at 408 nm and is indicative of heme-containing species.

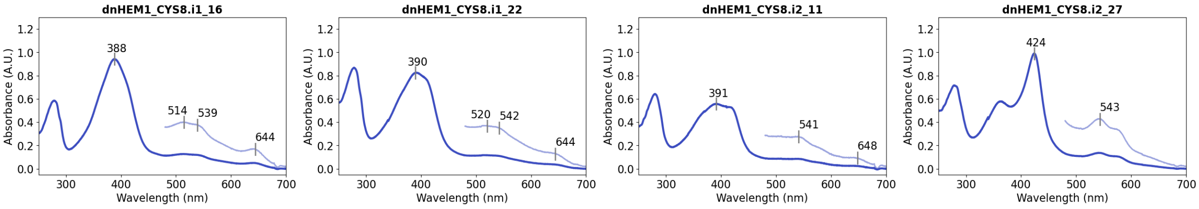

**Supplementary Figure 24.** UV-Vis spectra of heme-loaded inpainting-diversified dnHEM1_CYS8 variants. Inset shows the visible region at 3x magnification. Spectra were recorded in a buffer containing 25 mM Tris 300 mM NaCl at pH 8.

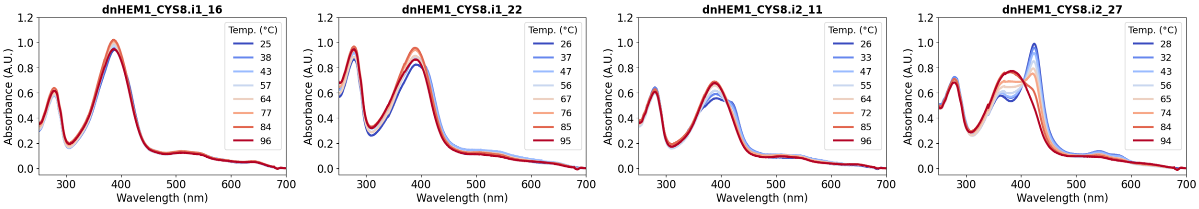

**Supplementary Figure 25.** UV-Vis spectra of heme-loaded inpainting-diversified dnHEM1_CYS8 variants collected at increasing temperatures (from room temperature to around 95 °C) at ~10 °C interval.

#### 3.5. Design of cysteine-ligated heme-binding proteins with RFdiffusion All-Atom

RFdiffusion All-Atom was used to generate a panel of cysteine-ligated heme-binding proteins. We have previously generated Cys-ligated heme-binding proteins based on RFdiffusion All-Atom protein backbone generation[2]. To expand this set, we started new design trajectories from a minimal active site model consisting of an axially cysteine-ligated heme with a spatial placeholder in the distal site (as was used in our previous work[2]). Sequences were generated using ProteinMPNN, and their structure predicted with AlphaFold2. Predicted structures of well-folding sequences were used as input in local heme binding site design using LigandMPNN-FastRelax. Finally, the obtained sequences were once again predicted with AlphaFold2, and successful variants further analysed by Rosetta (Supplementary Figure 26. Computational workflow for the design of CYS-ligated heme-binding proteins using RFdiffusion All-Atom, ProteinMPNN, LigandMPNN and AlphaFold2. Vertically stacked steps were performed in parallel.Supplementary Figure 26).

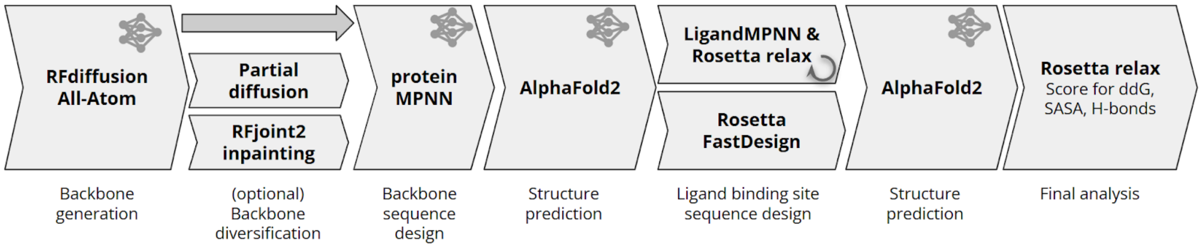

**Supplementary Figure 26.** Computational workflow for the design of CYS-ligated heme-binding proteins using RFdiffusion All-Atom, ProteinMPNN, LigandMPNN and AlphaFold2. Vertically stacked steps were performed in parallel.

**Experimental characterization**

Additional heme-binding proteins that were created using RFdiffusion All-Atom were produced and characterized using the methods described in our previous publication. Cloning and small scale (4 mL) *E. coli* culture preparation was performed following the “general protocol for cloning and testing designed proteins for heme binding”.

Proteins were isolated as clarified cell lysate, purified with Ni-NTA IMAC, 10 µM hemin was added to the protein-containing eluate which was then cleaned using a desalting column. UV-Vis spectroscopy was used to identify promising heme-binding proteins based on the presence of a significant Soret band in the 380-430 nm range. Cells producing promising candidates were isolated as clonal variants (verified by Sanger sequencing), the protein was produced on a larger scale (in a 40 mL TB-II culture), and purified in the presence of excess hemin. The oligomeric state, heme binding and thermal stability of these proteins was characterized using size exclusion chromatography, and UV-Vis spectroscopy, respectively (Supplementary Figure 27 & 28).

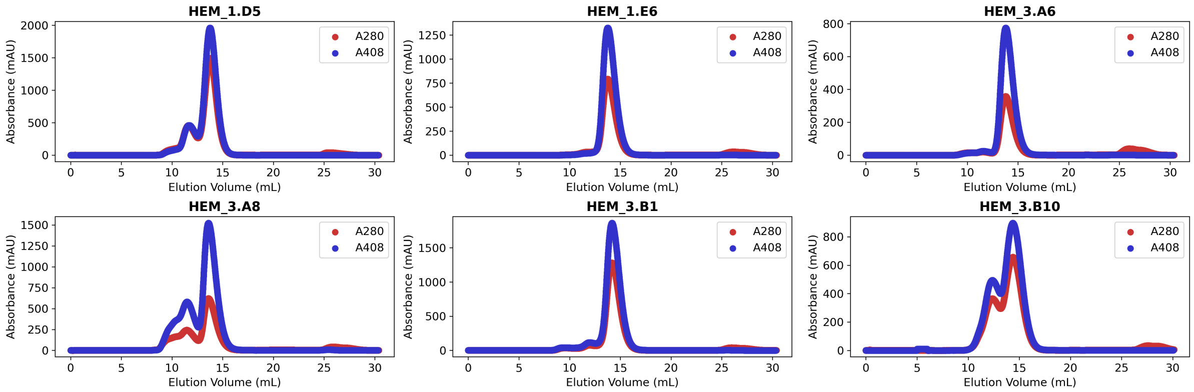

**Supplementary Figure 27.** Size-exclusion chromatograms of heme-loaded proteins. Data were collected using a Superdex Increase 75 10/300 GL column (GE Healthcare) in a buffer containing 50 mM KPi and 200 mM NaCl at pH 7.2. Void volume of the column is 8.5 mL. Blue chromatograms were obtained by following the absorbance at 408 nm, indicating elution of heme-containing species. Red chromatograms were obtained from absorbance at 280 nm.

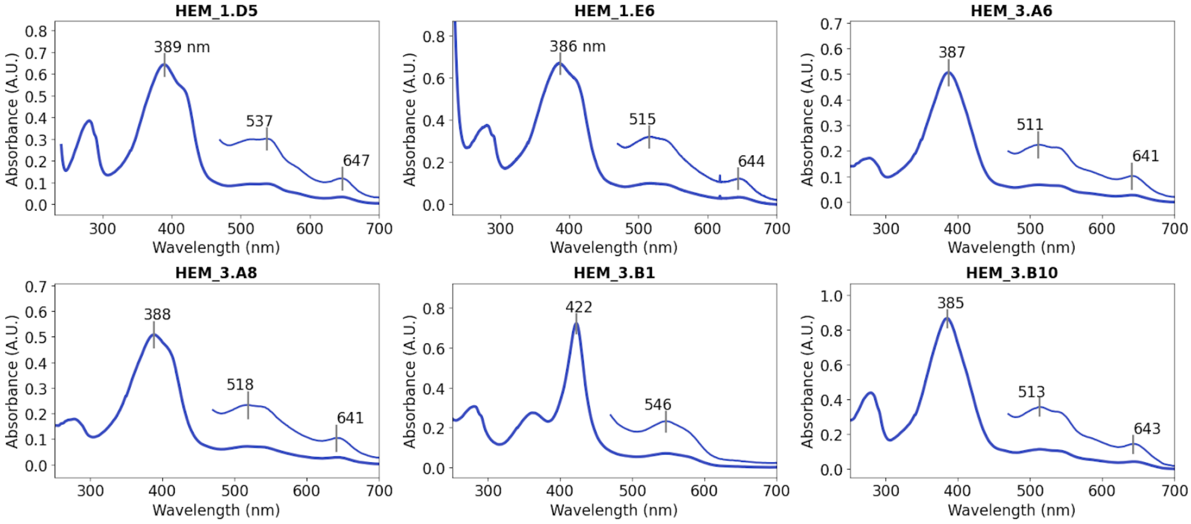

**Supplementary Figure 28.** UV-Vis spectra of heme-loaded proteins. Inset shows the visible region at 3x magnification. Spectroscopic data of most designs is in agreement with CYS-ligated heme binding (either Soret maximum at ~420 nm and Q band features at 540/570 nm for hexacoordinate low spin state, or Soret maximum at 370-390 nm and Q band features at 510/540 and charge transfer band at 640 nm for pentacoordinate high spin state). Spectra were recorded in a buffer containing 50 mM KPi and 200 mM NaCl at pH 7.2.
